# Tonic interferon restricts pathogenic IL-17-driven inflammatory disease via balancing the microbiome

**DOI:** 10.1101/2021.04.21.440756

**Authors:** Isabelle J. Marié, Lara Brambilla, Doua Azzouz, Ze Chen, Gisele Baracho, Azlann Arnett, Haiyan S. Li, Weiguo Liu, Luisa Cimmino, Pratip Chattopadhyay, Gregg Silverman, Stephanie S. Watowich, Bernard Khor, David E. Levy

## Abstract

Maintenance of immune homeostasis involves a synergistic relationship between the host and the microbiome. Canonical interferon (IFN) signaling controls responses to acute microbial infection, through engagement of the STAT1 transcription factor. However, the contribution of tonic levels of IFN to immune homeostasis in absence of acute infection remains largely unexplored. We report that STAT1 KO mice spontaneously developed an inflammatory disease marked by myeloid hyperplasia and splenic accumulation of hematopoietic stem cells. Moreover, these animals developed inflammatory bowel disease. Profiling gut bacteria revealed a profound dysbiosis in absence of tonic IFN signaling, which triggered expansion of T_H_17 cells and loss of splenic T_reg_ cells. Resolution of dysbiosis by antibiotic treatment averted the T_H_17 bias, and blocking IL17 signaling prevented myeloid expansion and splenic stem cell accumulation. Thus, tonic IFNs regulate gut microbial ecology, which is crucial for maintaining physiologic immune homeostasis and preventing inflammation.

## Introduction

Interferons (IFNs) are important mediators of innate and adaptive immunity. The generic term IFN comprises three types of cytokines: Type I family (IFN-I) encoded by multiple genes, including primarily numerous IFN-*α* subtypes and IFN-*β*; type II family (IFN-II), with IFN-*γ* being its sole member; and type III (IFN-III) family consisting of several IFN-*λ*s. Each IFN family signals through a distinct heterodimeric cell surface receptor. All members of IFN-I bind a receptor termed IFNAR, which triggers activation of the Jak kinases Jak1 and Tyk2 that mediate tyrosine phosphorylation of two members of the signal transducer and activator of transcription (STAT) family, STAT1 and STAT2. Activated STAT1 and STAT2, along with interferon regulatory factor (IRF) 9, form the heterotrimeric complex ISGF3 that binds to interferon stimulated response elements in the promoters of hundreds of interferon stimulated genes [1]. While IFN-III binds a distinct receptor composed of IL28Ra and IL10Rb subunits, this signaling cascade also activates ISGF3 and is largely overlapping with the pathway downstream of IFN-I [2]. In contrast, IFN-II, after binding its cognate receptor (IFNGR), signals predominately through homodimers of STAT1 and stimulates a set of genes containing a gamma-activating sequence (GAS) [3]. All these pathways converge on STAT1, and STAT1 deficiency or hypofunction leads to insensitivity to all types of IFN. As expected, human STAT1 deficiency results in increased susceptibility to both viral and mycobacterial infections [4, 5], and hematopoietic cell transplantation remains the only curative treatment [6]. Inability of STAT1-deficient patients to thrive in absence of mounting appropriate innate immune responses to microbes precludes study of a contribution of STAT1 in homeostasis. However, some STAT1 partial loss of function (LOF) patients suffer from chronic colitis, as well as severe infections [7, 8]. On the other hand, individuals with STAT1 gain-of-function (GOF) mutations suffer most frequently from mucocutaneous diseases, in part due to depressed levels of T_H_17 cells, thereby attributing important regulatory functions to STAT1 [9, 10].

Despite engaging similar downstream signaling cascades, it is becoming increasingly evident that IFN-I and –III play distinct roles in establishing innate immunity against microbes and participate differently in the overall immune functions of the host [11]. For instance, IFN-III has been shown to resolve inflammation by reducing the number of IL-17-producing T_H_17 helper T cells and restricting the recruitment of neutrophils [13]. Interestingly, besides its important role in inflammation, IL-17 also plays a significant role in hematopoiesis [14]. For instance, IL-17 stimulates myeloid and erythroid progenitors [15, 16], suggesting that IFN-III could play a role in the regulation of hematopoiesis through limiting the action of IL-17. Moreover, in part because of the more limited distribution of their receptor, IFN-III members display a unique capacity to counter pathogen invasion at mucosal sites while curbing overexuberant inflammation that helps maintain barrier integrity [12].

There is mounting evidence that commensal gut microbiota that exist at mucosal surfaces also play fundamental roles in shaping the host immune system [17–21]. However, there is only limited understanding of the importance of the interplay between the microbiome and IFN to prevent pathogenesis. Herein, we unravel a role for STAT1 in the control of microbiota ecology that prevents inflammation and maintains immune homeostasis in absence of an infectious challenge.

## Results

### STAT1 deficient mice exhibit splenomegaly, neutrophilia and increased splenic progenitors

We observed that STAT1 deficient animals developed a spontaneous splenomegaly irrespective of age and sex. STAT1 KO spleens were found to average 5 to 10 times larger than their wild type (WT) littermates (Fig. 1A) in absence of any notable infection. Moreover, the normal splenic architecture of STAT1 KO animals was disrupted, with prominent signs of extramedullary hematopoiesis (Fig. 1A). The number of total white blood cells (WBC), as well as neutrophil and monocyte subsets, was dramatically increased in STAT1 KO mice compared to WT (Fig. 1B and C).

**Fig. 1:**
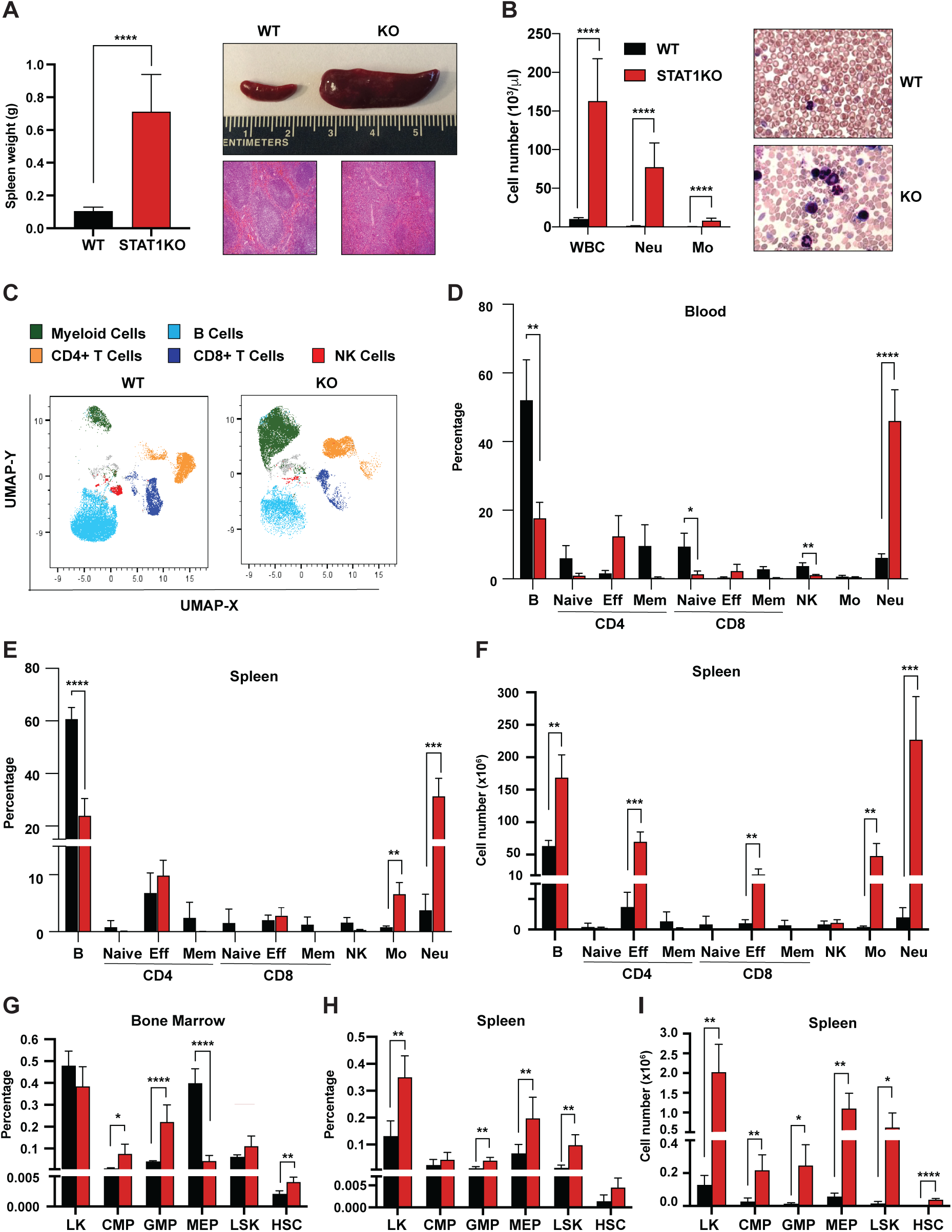
Characterization of STAT1 KO hematopoietic defects. A- Spleen weights in g (n = 7), mean ± S.D (left panel). Representative spleen of a STAT1 KO mouse and its age and sex matched WT littermate (top panel). Representative histology of STAT1 KO and WT spleens, stained with H&E (bottom panel) B- Total WBC, neutrophil and monocyte counts (n = 6 WT, n = 11 KO), mean ± S.D (left panel). Representative blood smear from WT and STAT1 KO blood after Giemsa staining (right panel). C- Uniform Manifold Approximation and Projection (UMAP) plots of blood from WT and STAT1 KO littermates after 18-color multi-parameter flow cytometry staining. Clusters were annotated using known markers. Flow cytometric analysis of blood (D), bone marrow (G), spleen (E-H) of STAT1 KO and WT littermates (n=4-5). Values represent mean ± S.D of live cells, percentage or number, as indicated. \**p<*0.05, ** *p<* 0.01, *** *p*< 0.001, **** *p<*0.0001 by student’s t-test. NK, natural killer; LK, lin^-^Sca1^-^c-Kit^+^; CMP, common myeloid progenitor; GMP, granulocyte- macrophage progenitor; MEP, megakaryocyte-erythroid progenitor; LSK, lin^-^Sca1^+^c-Kit^+^; HSC, hematopoietic stem cells.

To better characterize the profound alterations of the STAT1 KO hematopoietic system, we undertook a more comprehensive analysis of blood populations using multiparameter flow cytometry (Table S1, Fig. S1). WT and STAT1 KO blood were compared by visualization of Uniform Manifold Approximation and Projection (UMAP) plots. Profound changes in the distribution of the major classes of leukocytes were observed as shown in Fig. 1C. Consistent with the analysis of blood counts, a dramatic elevation in the percentage of monocytes and neutrophils was noted. The percentage of neutrophils increased from an average of 7% in WT animals to about 50% of total blood cells in STAT1 deficient mice (Fig. 1D). Conversely, STAT1 KO blood showed a compensatory loss of B lymphocytes and near disappearance of NK cells when compared to WT littermates. Interestingly, although the percentage of T cells did not change significantly, a notable shift from naïve and central memory to effector T cells was observed for both CD4^+^ and CD8^+^ populations (Fig. 1D). The same qualitative changes were observed when the analysis was performed on spleens, except that the percentage increase of effector T cells was less pronounced (Fig. 1E). Notably, because of the greatly increased cellularity of STAT1 KO spleens, the total number of monocytes and neutrophils was approximately 100-fold higher in the KO spleen compared to WT (Fig. 1F). Likewise, despite the differences in the percentage of effector T cells not being statistically significant, STAT1 KO spleens contained close to 10 times more CD4^+^ and CD8^+^ effector T cells as compared to WT spleens, and these quantitative differences were highly statistically significant (Fig. 1E and F).

The dramatic changes observed in the populations of mature cells in blood and spleen prompted us to examine stem and progenitor cells in bone marrow and spleen. Analysis of progenitor populations in the bone marrow showed significant increases in GMP accompanied by a decrease of MEP in STAT1 KO compared to WT bone marrow (Fig. 1G). In contrast, we observed an increase of MEP in the spleens of STAT1 KO mice, consistent with the presence of extramedullary hematopoiesis. More strikingly, the number of phenotypic hematopoietic stem cells (HSCs), defined as Lin^-^Sca1^+^c-Kit^+^CD150^hi^CD48^low^, which is very low in WT spleen, was significantly expanded in STAT1 deficient spleens (Fig. 1H and I).

### Definitive HSCs populate STAT1 KO spleens

In order to confirm that the Lin^-^Sca^+^Kit^+^CD150^hi^CD48^low^ cells present in STAT1 KO spleens were functional HSCs, we first tested their self-renewing potential *in vitro*. We compared the plating efficiency in myeloid promoting methylcellulose cultures of splenic and bone marrow cells from WT and STAT1 KO animals. Plating efficiency of STAT1 deficient bone marrow cells was higher than WT for the first plating, but subsequent passages showed only minor differences between the 2 genotypes (Fig. 2A). In contrast, STAT1 deficient splenic progenitors showed much higher plating efficiency compared to WT, as well as greatly enhanced replating capacity. Indeed, STAT1 deficient splenic progenitors still gave rise to a significant number of colonies after four sequential passages in methylcellulose (Fig. 2B). This result provides evidence that STAT1 deficient spleens host a large population of self-renewing progenitors that is not normally found in WT animals.

**Fig. 2:**
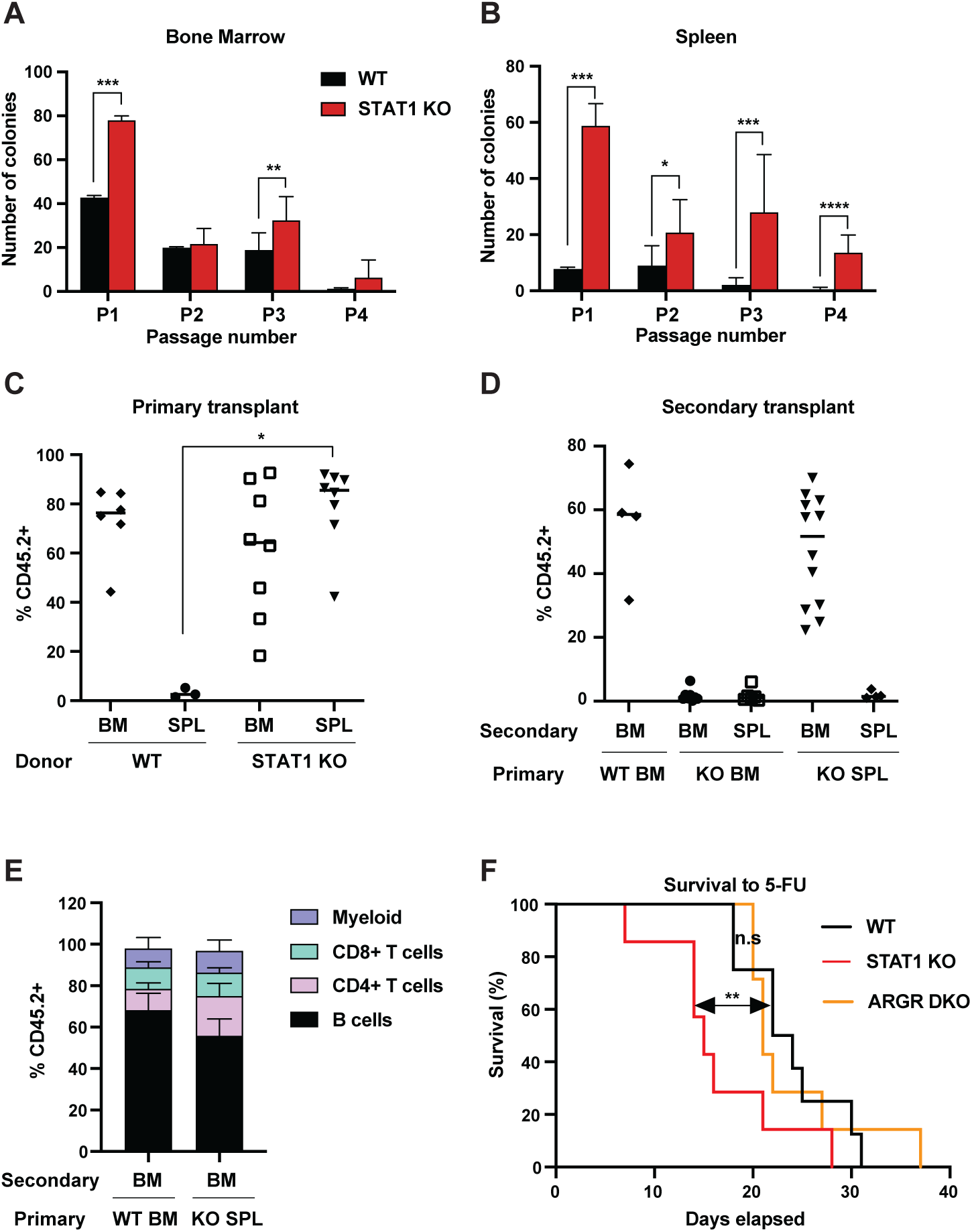
STAT1 deficient spleens harbor a large number of definitive HSCs. In vitro self-renewal colony forming assay of hematopoietic progenitors from bone marrow (A) or spleen (B) after serial plating (n=3 mice, each plating in triplicate). Values represent mean number of colonies ± S.D. Each replating culture was inoculated with equal numbers of cells. Quantification of donor-derived (CD45.2) cells, provenance as indicated, in the peripheral blood of recipient CD45.1 animals 1 mo after primary transplant (C) or 4 mo after secondary transplant (D). Values represent individual animals. Quantification of donor-derived (CD45.2) B (B220^+^) cells, T (CD4^+^ or CD8^+^), and myeloid (CD11b^+^) cells in the peripheral blood of recipient animals 4 mo after secondary transplant (E); Values represent mean ± S.D.; WT n=4 mice; KO n=11 mice F- Kaplan-Meier survival curve of WT, STAT1 KO and ARGR DKO mice after weekly injections of 5- fluorouracil (5-FU). WT n=5 mice, STAT1 KO and ARGR DKO n=7 mice. * *p<*0.05, ** *p<* 0.01, *** *p<* 0.001, **** *p<*0.0001 by student’s t-test.

Since self-renewal of hematopoietic cells *in vitro* largely scores the presence of short-term progenitors, we wanted to assess the presence of functional long-term (LT) HSCs in STAT1 deficient spleens. Therefore, we performed a transplantation experiment into lethally-irradiated recipient animals (CD45.1) using both spleen and bone marrow of WT and STAT1 KO animals (CD45.2) as donor cells. Reconstitution of the hematopoietic system by donor cells was monitored after 1 mo. Strikingly, while WT splenic cells were not capable of thriving in irradiated hosts, indicative of the limited presence of HSCs in this organ under normal conditions [22], STAT1 deficient splenic cells reconstituted the host hematopoietic system as efficiently as WT or STAT1 KO BM, with a median reconstitution of over 80% donor-derived cells (Fig. 2C). We next harvested the bone marrow and spleen from primary engrafted recipients and transplanted these cells into irradiated secondary recipients (CD45.1), in order to distinguish between short-term and long-term engraftment capability. Results from the secondary transplantation experiments showed even greater differences between WT and KO cells. Neither bone marrow nor spleen cells isolated from primary reconstituted animals that had received STAT1 KO bone marrow were capable of complementing secondary transplanted recipients, suggesting that STAT1 KO bone marrow was deficient in LT- HSCs (Fig. 2D). In marked contrast, bone marrow cells from primary animals reconstituted with STAT1 KO spleen cells had a reconstituting capacity indistinguishable from animals initially reconstituted with WT bone marrow (Fig. 2D). Finally, in order to confirm the pluripotency of the progenitors found in STAT1 deficient spleens, blood from secondary transplanted recipients was analyzed after 4 mo. We found no significant differences of the lineage allocation of blood amongst secondary recipients (Fig. 2E). These results strongly suggest that fully functional pluripotent LT-HSCs reside in STAT1 deficient spleens, that these cells, albeit residing in the spleen of STAT1 deficient animals, home to the BM following engraftment of WT animals, and that STAT1 KO BM is either depleted of LT-HSCs or these cells are functionally deficient. Of note, no myeloid expansion or splenomegaly was detected in animals transplanted with STAT1 KO bone marrow or spleen, suggesting that the myeloproliferative disease observed in STAT1 KO mice is not fully cell intrinsic.

We considered whether the increased abundance of splenic LT-HSCs present in STAT1 KO animals represented an altered proliferative capacity. To investigate this possibility, we tested the sensitivity of STAT1 KO mice to 5-fluorouracil (5-FU), a myelosuppressive chemotherapeutic that kills proliferating cells. Following weekly injections of 5-FU, we documented that STAT1 KO mice died significantly faster than WT, and also faster than mice lacking both IFNAR and IFNGR (ARGR DKO) (Fig. 2F). Since the lethality of 5-FU is related to depletion of cycling stem and progenitor cells, this result suggests that the increased abundance of HSCs observed in STAT1 KO spleen are more sensitive to depletion than WT stem cells, perhaps due to decreased quiescence or more rapid exhaustion following cycling. Increased progenitor proliferation would be consistent with the enhanced replating efficiency of these cells observed *in vitro* (Fig. 2B). Notably, the response of ARGR DKO animals to 5-FU challenge was indistinguishable from WT animals, suggesting that IFN-I and IFN-II are not required to regulate HSC quiescence, at least in the presence of IFN-III.

### STAT1 deficient CD4 T cells produce elevated levels of IFN-γ and IL-17A

In order to better understand the underlying mechanisms prompting the inflammatory phenotype observed in a STAT1 deficient hematopoietic system, we undertook a comprehensive cytokine and chemokine profiling of the sera of these animals (Data File S1). IFN-*γ* was increased approximately 300-fold in the sera of STAT1 KO animals and was moderately elevated in ARGR DKO mice (Fig. 3A). IL-5 and TNF-*α* showed a similar profile but reduced magnitude relative to IFN-*γ* (Fig. 3A). However, a more striking difference was noted for IL-17A, which was found to average 100-fold higher concentration in STAT1 KO sera, with little or no elevation in ARGR DKO (Fig. 3A). Since ARGR DKO animals, despite their lack of responsiveness to IFN-I and –II, showed little to no sign of pathology, the high concentrations of circulating IL-17A correlated with disease.

**Fig. 3:**
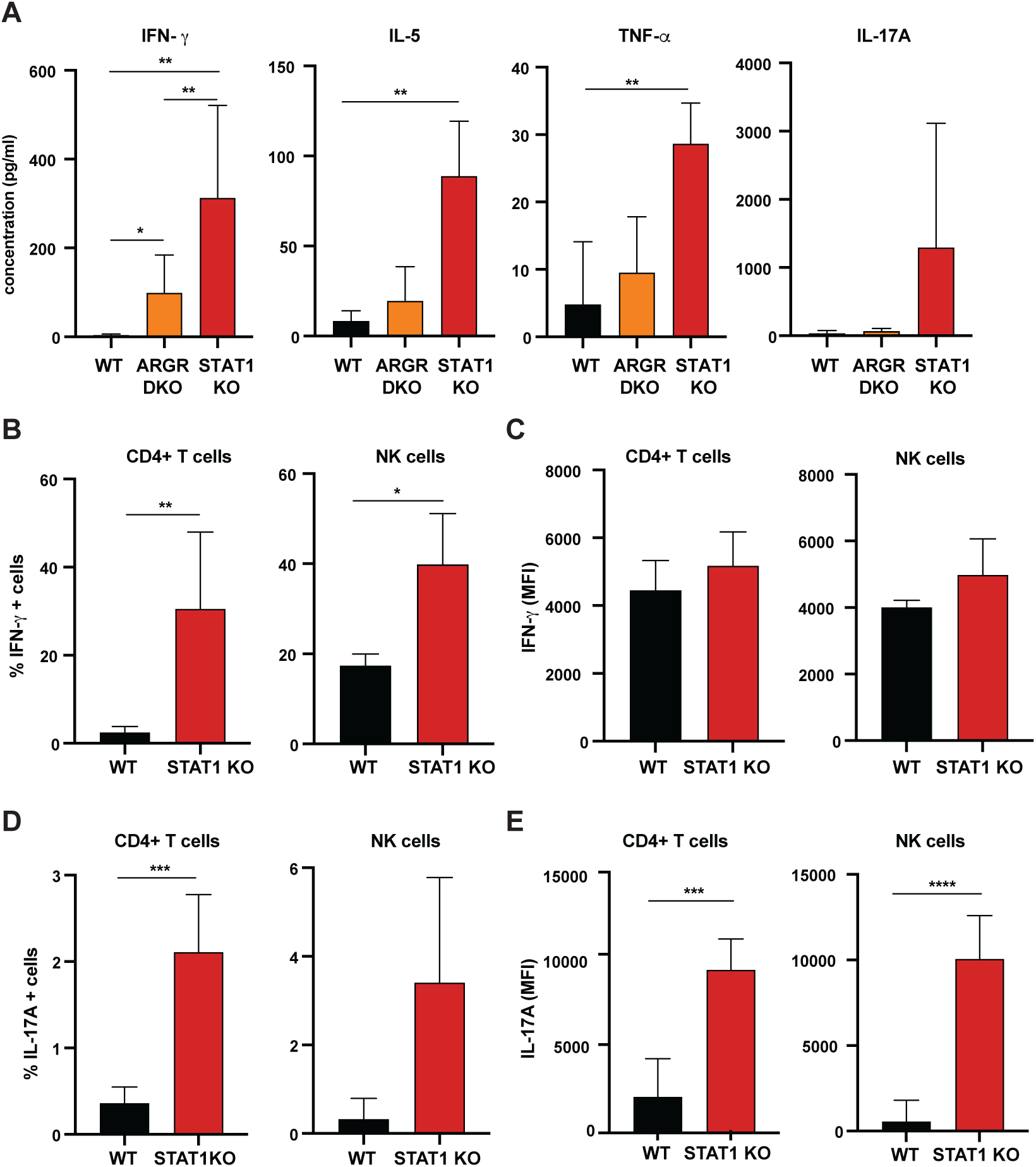
Cytokine profiling of STAT1 KO blood. A- Cytokine concentration in blood of WT, ARGR DKO and STAT1 KO mice in pg/ml (n=6-9 mice for each group). Data are representative of two experiments. Percentage of IFN-γ (B) or IL-17A-producing (C) CD4+ T cells (left panel) or NK cells (right panel) in STAT1 KO and WT mice (n=5 or 6) with corresponding MFI (D-E). Values represent mean ± S.D. * *p<*0.05, ** *p<* 0.01, *** *p<* 0.001, **** *p<*0.0001 by student’s t-test.

Given the observation of elevated cytokine levels, we sought to determine what cells were the source of IFN-*γ* and IL-17A. Therefore, we assayed peripheral blood lymphocytes by intracellular cytokine flow cytometry. We found that CD4^+^ T cells and NK cells were the main populations making IFN-*γ* and IL-17A. As many as 30% of STAT1 deficient CD4^+^ T cells produced IFN-*γ* as compared to only a few percent in WT controls (Fig. 3B, C). Similarly, a larger percentage of STAT1 KO CD4^+^ T and NK cells made IL- 17A when compared to WT counterparts (Fig. 3D). Moreover, not only was the number of IL-17 producing cells greater in STAT1 KO animals, but the amount of cytokine produced per cell was much greater, as indicated by a 5-10-fold increased mean fluorescence intensity (Fig. 3E).

### STAT1 KO pathology is associated with combined insensitivity to IFN-I, -II and –III

STAT1 is a major transcription factor mediating the canonical JAK-STAT pathways downstream all three types of IFN. However, IFNs can also activate accessory pathways that could become preponderant in absence of STAT1 and trigger an imbalance of the hematopoietic system homeostasis [23]. To further evaluate a role for IFN, we applied a genetic approach and bred our STAT1 KO colony with mouse strains deficient for IFNAR and IFNGR that renders them insensitive to type I and II IFNs. The resulting triple deficient strain that lacked IFN-I and IFN-II receptors (ARGR STAT1 TKO) still displayed the same altered parameters that we observed in mice singly deficient for STAT1 (Fig. S2). Therefore, alternate signals downstream of IFN receptors are unlikely culprits of the observed pathology. However, loss of IFN-I and II signaling (ARGR DKO) did not mimic the phenotype mediated by STAT1 loss. These data suggested that STAT1 was essential to immune fitness but the role of IFNs remained unclear.

To further investigate roles for IFNs upstream of STAT1 in providing protection from hematopoietic dysregulation, we examined the phenotypes of mice unable to respond to IFN-I, -II, and –III. As previously noted, ARGR DKO mice showed very mild signs of an altered immune system compared to STAT1 KO animals. Despite being unresponsive to IFN-I and –II, these animals did not present with splenomegaly, and displayed very mild neutrophilia (Fig.4A). Interestingly, these animals did not exhibit an increase of CD4^+^ effector T cells as observed in the STAT1 KO blood and spleen (Fig. 4B). In contrast, STAT2 KO animals, that are impaired in their response to IFN-I and –III, showed profound defects that largely recapitulated what was observed in STAT1 deficient mice, except that no increase in the percentage of CD8+ effector T cells was detected in blood (Fig. 4 B, C, D). Taken together, these data suggested that IFN-I and –III are the main cytokines acting through STAT1 to provide a protective environment to insure proper homeostasis of the immune system, but additional STAT1-dependent functions are also at play. To better examine a role for type II IFN in this process, we compared STAT2 KO and IFNGR/STAT2 (GR STAT2) DKO mice. GR STAT2 DKO animals cannot respond to any type of IFN, similar to the IFN signaling defect in STAT1 KO mice. This strain also provided the advantage of probing any role for additional cytokines or growth factors that activated STAT1 but not STAT2. Detailed flow cytometry analysis of the blood and spleen of GR STAT2 DKO mice uncovered defects virtually identical to the ones observed for STAT1 KO mice (Fig. 4B and C), strongly indicating that the phenotype of STAT1 deficient mice is associated with combined insensitivity to IFN-I, -II and –III (Table S2).

**Fig. 4:**
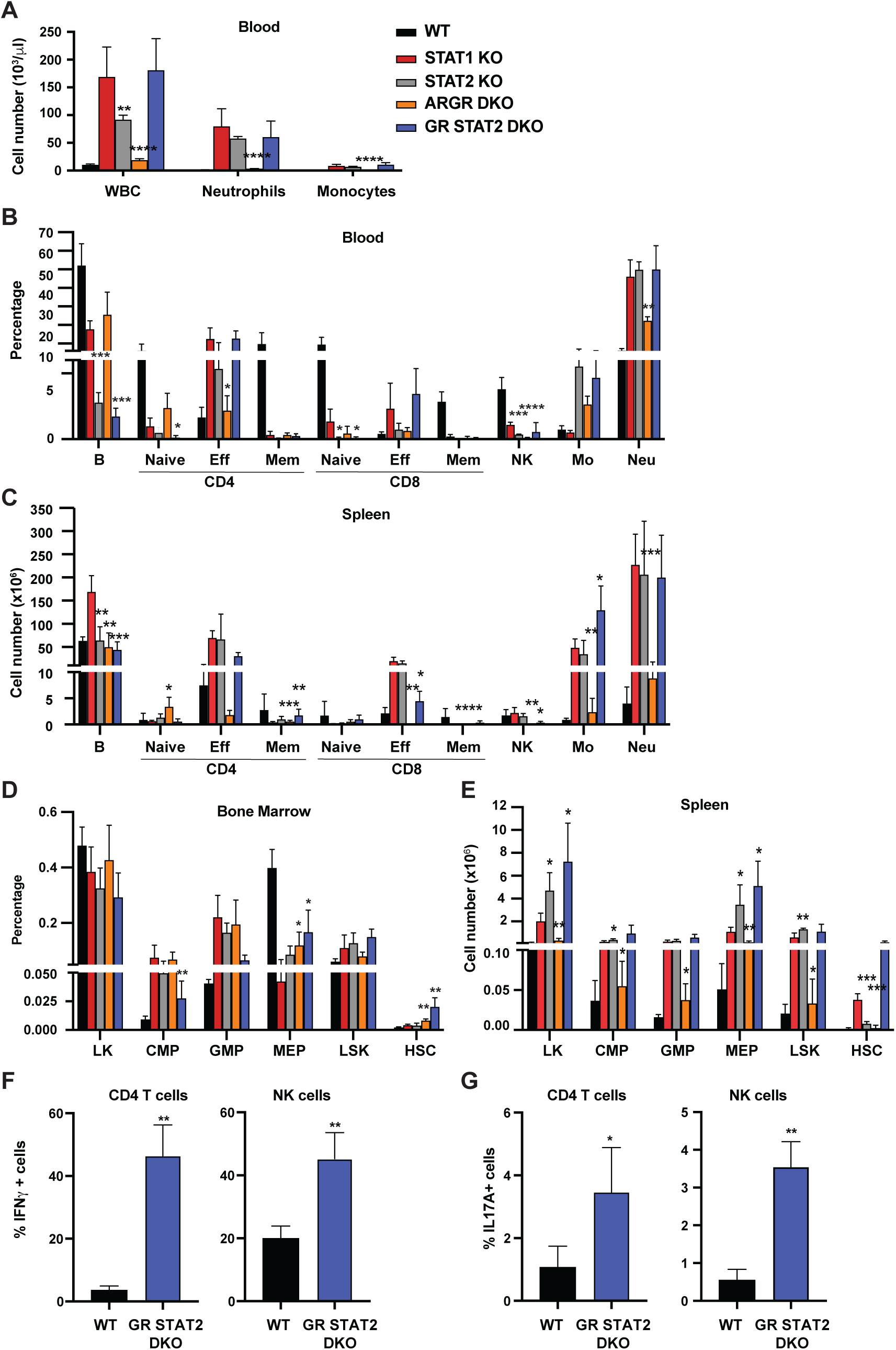
Combined IFN type I, II, III response deficiency recapitulates STAT1 deficiency. A- Total WBC, neutrophils and monocytes counts using Hemavet in blood from WT, STAT1 KO, STAT2 KO, ARGR DKO and GR STAT2 DKO mice (n=5-10 mice/group). Quantification by flow cytometry of mature cell populations from blood (B), spleen (C), and progenitor populations from bone marrow (D) and spleen (E) from WT, STAT1 KO, STAT2 KO, ARGR DKO and GRSTAT2 DKO (n=4 or 5). Values represent mean ± S.D of percentage of single live cells, or cell number for spleen. Same legend as in A. Statistical comparison is between each genotype and STAT1 KO. Percentage of IFN-γ (F) or IL-17A-producing (G) CD4+ T cells (left panel) or NK cells (right panel) in GR STAT2 DKO and WT mice (n=4-6). Values represent mean ± S.D of live cells, percentage or number, as indicated. * *p<*0.05, ** *p<* 0.01, *** *p<* 0.001, **** *p<*0.0001 by student’s t-test. NK, natural killer; LK, lin^-^Sca1^-^c-Kit^+^; CMP, common myeloid progenitor; GMP, granulocyte-macrophage progenitor; MEP, megakaryocyte-erythroid progenitor; LSK, lin^-^Sca1^+^c-Kit^+^; HSC, hematopoietic stem cells.

Interestingly, analysis of bone marrow progenitors showed an increase of CMP and GMP accompanied by a decrease of MEP representation for all mutant genotypes studied (Fig. 4D). More importantly, the number of splenic HSCs was dramatically increased in STAT1 KO and GR STAT2 DKO mice, and this increase was not observed in the other mutant strains (Fig.4E). Taken together, our results strongly suggested that the combined absence of IFN-*γ*R and STAT2 (GR STAT2 DKO) phenocopied STAT1 deficiency. To confirm this hypothesis, we measured IFN-*γ* and IL-17A production by CD4^+^ T cells and NK cells in these animals. Both cytokines were elevated compared to WT (Fig. 4F and G), mimicking the phenotype of STAT1 KO cells (Fig. 3).

### STAT1-deficient mice are prone to develop colitis

In addition to the hematopoietic defects documented above, STAT1 KO mice developed spontaneous colitis, frequently accompanied by rectal prolapse. Gross anatomy of the STAT1 deficient colons revealed increased angiogenesis (Fig 5A), a characteristic of inflamed bowels [24]. Histopathologic analysis of colon sections documented extensive immune cell infiltrates, epithelial hyperplasia and goblet cell loss in STAT1 KO colons, associated in some cases with the presence of cryptitis (Fig. 5 B and C). Damage to the gut epithelium often results in alteration of the intestinal barrier integrity and bacterial leakage into neighboring organs. Therefore, we scored for presence of bacteria in the liver, mesenteric lymph nodes and spleens of STAT1 KO mice compared to WT animals. Bacteria were observed mainly in the liver in STAT1 KO mice and to a lesser extent in the mesenteric lymph nodes and very rarely in the spleen (Fig. 5D), while organs from WT animals were largely sterile. In addition, we tested the sensitivity of STAT1 KO mice to dextran sulfate sodium (DSS)-induced colitis. Signs of DSS-induced intestinal inflammation, as evidenced by colon shortening and altered histopathology, were more pronounced in STAT1 deficient mice than WT littermates (Fig. 5E and F).

**Fig. 5:**
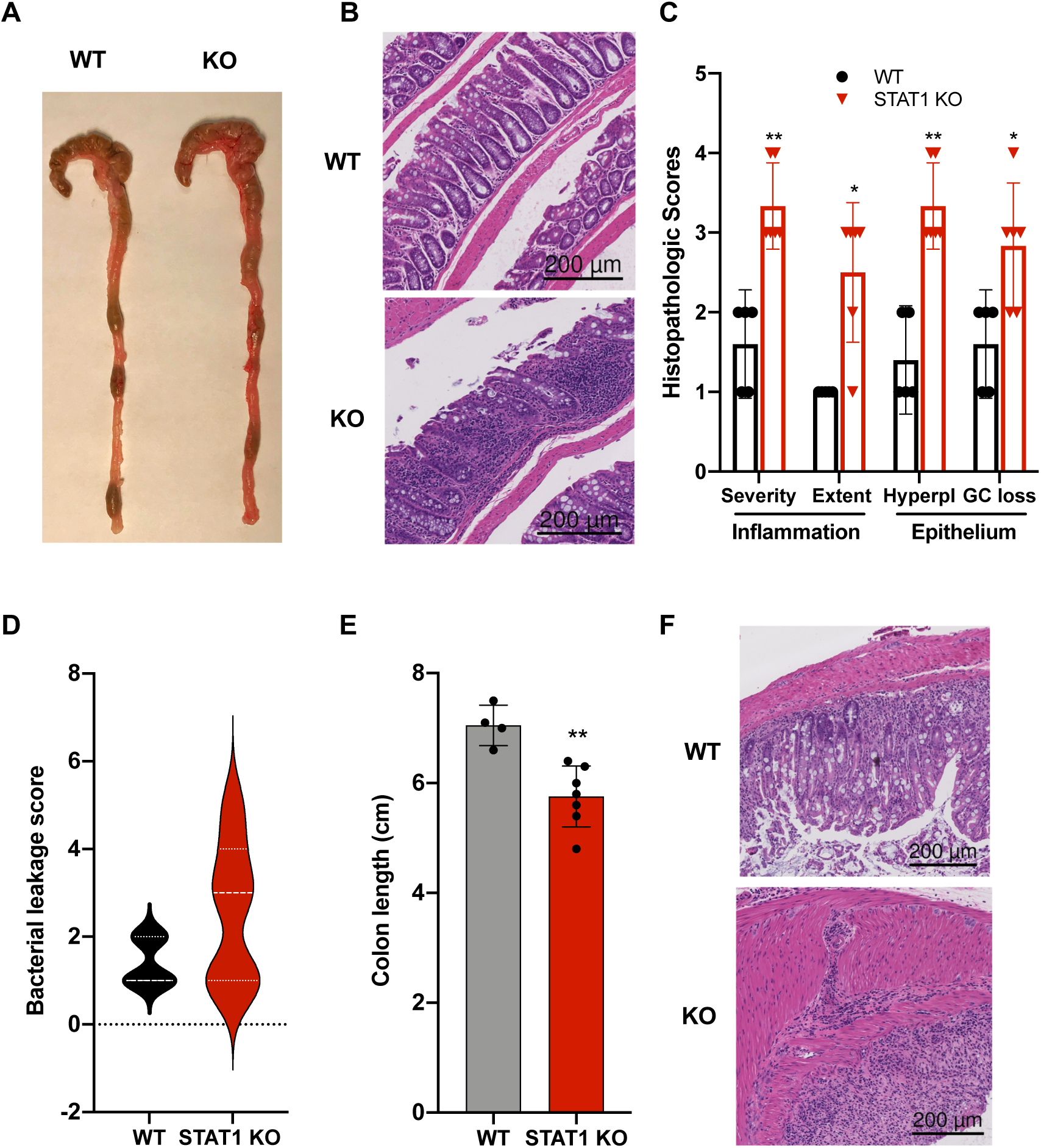
STAT1KO mice are prone to develop spontaneous colitis. A- Representative colon gross anatomy of WT and STAT1 KO mice Representative histology of WT and STAT1 KO colons, stained with H&E (scale bar = 200μm). B- Histopathologic scores of WT and STAT1 KO colons. Scoring was performed for severity and extend of inflammatory cell infiltrates and for epithelial changes (hyperplasia (hyperpl) and goblet cell loss (GC loss)). (n=5 for WT, n=6 for STAT1 KO) \**p<*0.05, ** *p<* 0.01 by Mann-Whitney *U* test. C- Bacterial leakage scores into liver, mesenteric lymph nodes and spleen as described in materials and methods (n=5 for WT, n=7 for STAT1 KO). D- Colon length after 7d of DSS treatment in drinking water; (n=4 for WT, n=7 for STAT1 KO) ** *p<* 0.01by student’s t-test. E- Representative histology of WT and STAT1 KO colons, stained with H&E, after 7d of DSS treatment (scale bar = 200μm).

### STAT1-deficient mice present with altered gut microbiota

Inflammatory bowel diseases (IBD) such as colitis are often associated with altered microbiota. Therefore, we investigated whether STAT1 deficiency could perturb the representation of gut microflora and cause dysbiosis. To this end, we performed 16S rRNA gene sequencing on fecal DNA samples collected from age- and gender-matched animals from 4 different colonies: WT, STAT1 KO, ARGR DKO and GR STAT2 DKO. Analysis of alpha-diversity measured by the Simpson and Shannon diversity indexes revealed a significant reduction of bacterial taxa diversity in STAT1 KO and GR STAT2 DKO animals compared to WT, whereas the ARGR DKO microbiome was found to be as diverse as the controls (Fig. 6A and B). Thus, commensal bacterial diversity was inversely correlated with the severity of the inflammatory disease. Furthermore, beta-diversity represented by principal coordinate analysis (PCoA) underscored similarities in bacterial diversity between the different groups of animals partially or completely lacking IFN response (Fig. S3A). Accordingly, analysis of the relative frequency of major bacterial families unveiled several significant differences between WT and mutant strains (Fig. 6C), among them a large decrease of *Bacteroidales S24-7* and *Burkholderiales Alcaligenaceae* and a consistent increase of *Deferribacterales Deferribacteraceae* and *Campylobacterales Helicobacteraceae*, suggesting that the abundance of these species is controlled by IFN (Fig. 6D). However, another family, *Bacteroidales Prevotellaceae*, was of particular interest, since increases were present only in STAT1 KO and GR STAT2 DKO animals, therefore correlating with disease severity (Fig. 6E). Two predominant clusters of amplicon sequence variants (ASV) related to the *Prevotella* family were increased in diseased animals. Since these ASV had not been taxonomically assigned to any previously identified species, we generated a phylogenetic tree based on the 16S rRNA sequences of these ASV and matched with known *Prevotella* taxa assignments (Fig. S3B). This analysis revealed that these ASV clustered with *Prevotella heparinolytica*, a microbial species known to trigger T_H_17 responses [25].

**Fig. 6:**
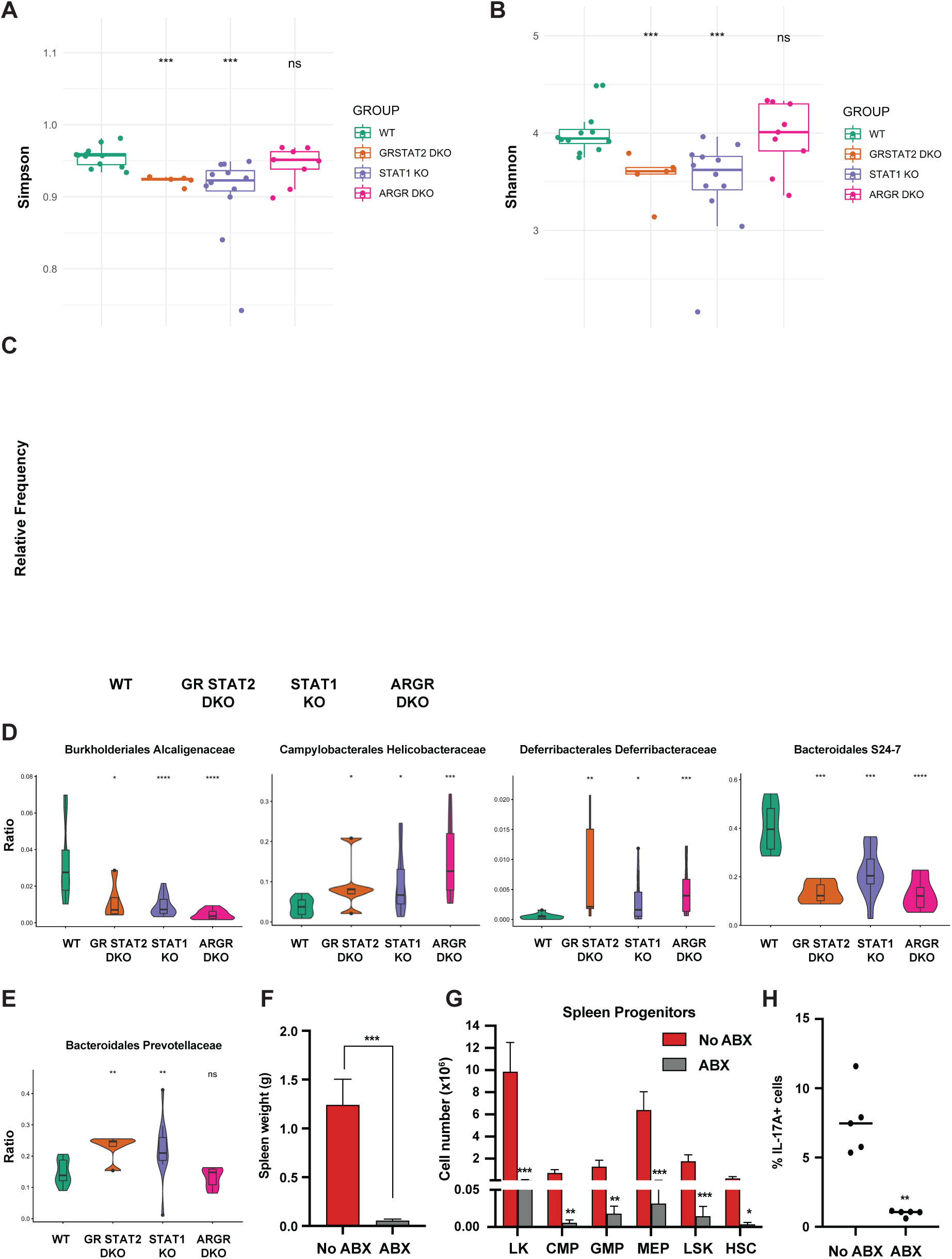
Tonic IFN controls gut microbiota. 16S rRNA sequencing of fecal DNA samples from WT, GR STAT2 DKO, STAT1 KO and ARGR DKO mice (n = 8-12 animals/strain). A- Alpha diversity as estimated by Simpson index (Kruskal-Wallis p value=0.00068) B- Alpha diversity as estimated by Shannon index (Kruskal-Wallis p value=0.00082) C- Family level microbiota composition D- Relative abundance of the total microbial load of four families of bacteria. E- Relative abundance of the total microbial load of Bacteroidales Prevotellaceae. F- Spleen weights in g (n= 4). G- Flow cytometric analysis of splenic progenitors of antibiotics (ABX)-treated STAT1 KO animals compared to untreated animals (No ABX) (n=4). H- Percentage of IL-17A-producing CD4+ T cells in antibiotics (ABX)-treated animals compared to untreated animals (No ABX). Values represent mean ± S.D of live cells, percentage or number, as indicated. *p<0.05, ** p< 0.01, *** p< 0.001, **** p<0.0001 by student’s t-test. LK, lin-Sca1-c-Kit+; CMP, common myeloid progenitor; GMP, granulocyte-macrophage progenitor; MEP, megakaryocyte-erythroid progenitor; LSK, lin-Sca1+c-Kit+; HSC, hematopoietic stem cells.

To probe for a possible causative connection between the observed dysbiosis of STAT1 KO animals and disease, we treated STAT1 KO mice with a mixture of broad- spectrum antibiotics (ABX). Following 4 wk treatment, analysis of fecal DNA samples from ABX-treated mice showed a greater than 100-fold reduction in total bacterial DNA, consistent with an expected depletion of commensal microbiota. Strikingly, ABX-treated STAT1 KO mice did not display splenomegaly (Fig. 6F). Moreover, ABX treatment restricted the accumulation of splenic CMP, GMP and MEP progenitors as well as HSCs in STAT1 KO animals (Fig. 6G). Finally, peripheral blood leukocytes from ABX-treated and control STAT1 KO mice were examined for IL-17A production. The elevated IL-17^+^ CD4^+^ T cells characteristic of STAT1 KO mice were significantly decreased by ABX treatment (Fig. 6H).

### IL-17Ra deficiency rescues splenomegaly and neutrophilia observed in STAT1 deficient mice

Inflammation in STAT1 KO mice strongly correlated with IL17 production and altered peripheral T cell profiles. To examine mechanisms underlying the T cell alterations, we first probed the ability of STAT1 KO CD4^+^ T cells to differentiate into T_H_17 cells *in vitro*. No difference in the generation of T_H_17 cells *in vitro* was observed between STAT1 KO and WT controls in response to IL-6 and TGF-*β* (Fig. S4A). We next analyzed the capacity of STAT1 KO CD4^+^ T cells to differentiate into T_reg_ cells *in vitro* in response to TGF-*β* and IL-2. Again, no significant deficit was observed in production of STAT1 KO T_reg_ compared to WT (Fig. S4B). Likewise, analysis of T cell populations in the thymus uncovered no statistically significant differences in thymic T_reg_ numbers (Fig. S4C). However, contrasting with what we observed *in vitro* and in the thymus, peripheral T_reg_ were virtually absent in STAT1 KO spleens (Fig. 7A), consistent with colitis [26] and with an inability to suppress T_H_17 expansion.

**Fig 7:**
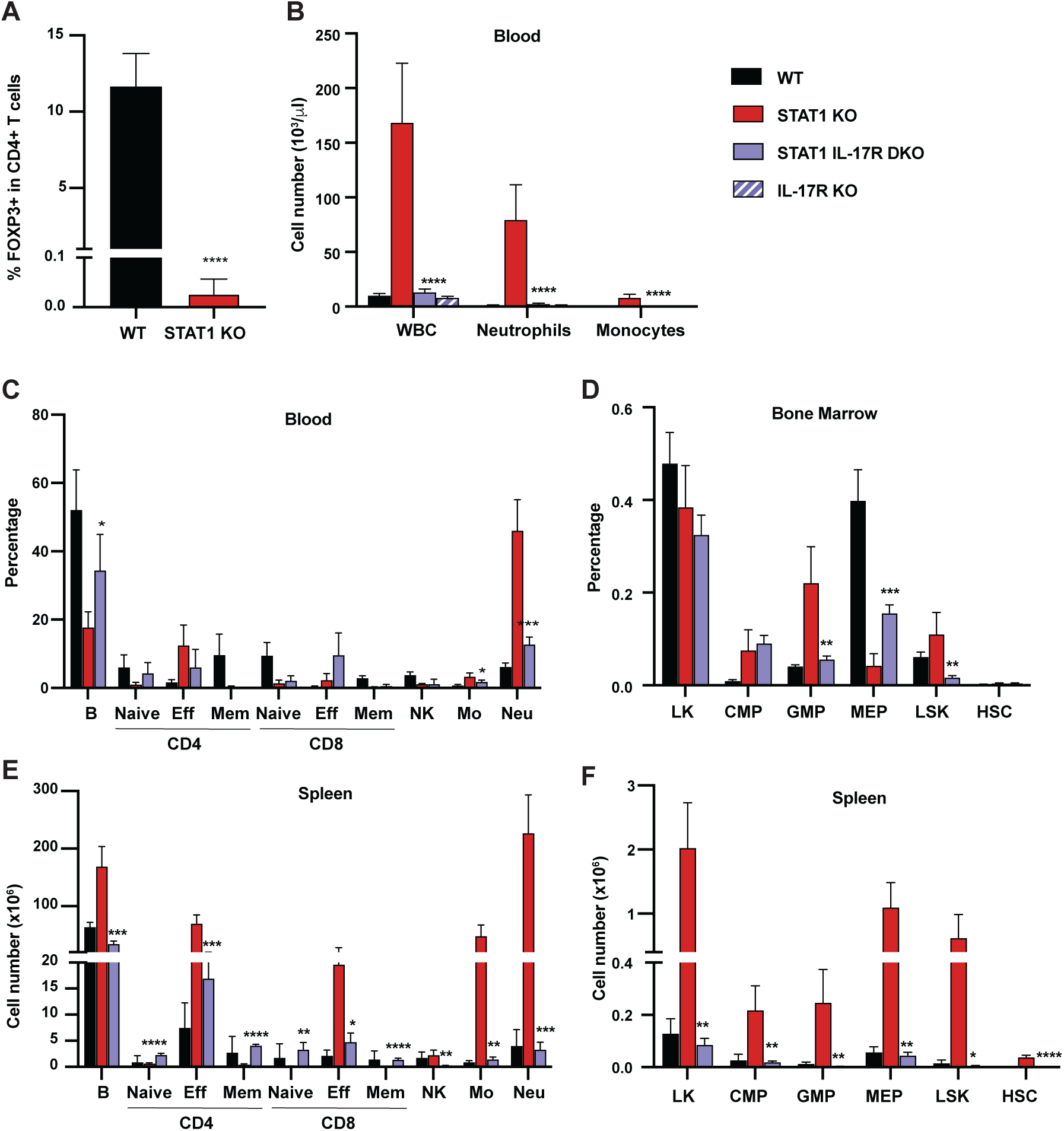
IL-17R deficiency rescues hematopoietic anomalies of STAT1 KO mice. A- Percentage of FOXP3^+^ T_reg_ cells in splenic CD4^+^ T cells of WT and STAT1 KO littermates (n=4). B- Total WBC, neutrophil and monocyte counts of WT STAT1 KO, IL-17R STAT1 DKO and IL- 17R KO, mean± S.D (n= 6 for WT, n=9 or 10 for other groups). Quantification by flow cytometry of mature cell populations from blood (C), spleen (F), and progenitor populations from bone marrow (D) and spleen (E) from WT, STAT1 KO, IL-17R STAT1 DKO (n=4 or 5). Values represent mean ± S.D of live cells, percentage or number, as indicated. \**p<*0.05, ** *p<* 0.01, *** *p<* 0.001, **** *p<*0.0001 by student’s t-test. NK, natural killer; LK, lin^-^Sca1^-^c-Kit^+^; CMP, common myeloid progenitor; GMP, granulocyte- macrophage progenitor; MEP, megakaryocyte-erythroid progenitor; LSK, lin^-^Sca1^+^c-Kit^+^; HSC, hematopoietic stem cells.

To directly test whether IL-17 could drive the anomalies observed in STAT1 deficient animals, we generated a mouse strain deficient for both STAT1 and IL-17Ra (STAT1 IL-17R DKO). Strikingly, blocking IL-17 responses normalized the blood parameters of STAT1 deficient mice. As expected, IL-17Ra single deficient mice did not present any sign of perturbed hematopoiesis, as previously described [27] (Fig. 7B). An in depth analysis of the representation of mature cells in the blood and spleen of these animals showed that STAT1 IL-17R DKO mice displayed at most a modest increase of the myeloid population (Fig. 7C and E). Likewise, concomitant IL-17Ra deficiency lowered the percentage of GMPs observed in STAT1 deficient bone marrow and spleen, resolving this phenotype essentially to control levels (Fig. 7D and F). More importantly, we observed no increase of splenic HSCs in STAT1 IL-17R DKO mice (Fig. 7F). As expected, production of IFN-*γ* and IL-17A by STAT1 IL-17R DKO CD4^+^ T cells and NK cells was not significantly different from that of STAT1 KO cells (Fig. S5A-D). Accordingly, the percentage of circulating effector T cells was equivalent to that seen in STAT1 KO mice (Fig. 7C). Taken together, these results point to increased IL-17A signaling as the primary driver of hematopoietic phenotypes in STAT1 KO animals but is downstream of alterations in lymphocyte subsets.

## Discussion

Our data provide evidence that tonic signaling through the IFN pathway, acting through its common mediator STAT1, is an essential modulator of immune homeostasis, reducing the propensity for inflammation. This tonic IFN signaling appears to involve all of the three major IFN families, since loss of all three arms of the pathway were required for the full inflammatory phenotype. Absence of tonic IFN signaling allows development of a skewed microbiome that triggers an inflammatory response, likely due to an increased bias of T_H_17 cells towards a pathogenic phenotype and impaired homing or survival of peripheral T_reg_.

A striking attribute of the inflammatory phenotype that developed in absence of STAT1 was a profound myeloid hyperplasia, accompanied by the accumulation of stem and progenitor cells in peripheral lymphoid organs. This phenotype was fully recapitulated in GR STAT2 DKO mice, demonstrating that it is caused by loss of IFN signaling, rather than by a non-canonical function of STAT1, and it was fully ameliorated by loss of IL17Ra, showing that IL17 signaling is epistatic to STAT1 for this phenotype. Myeloid hyperplasia in absence of STAT1 is not unprecedented, since a distinct strain of STAT1 KO mice housed in an independent animal facility was previously reported to develop a myeloid proliferative disease accompanied by progenitor expansion [28]. The fact that an independent strain of mutant mice housed in a distinct facility developed the same myeloid proliferative disease reinforces the notion that loss of signaling through the STAT1 pathway is a fundamental cause.

Most of the inflammatory phenotype that develops in absence of tonic IFN signaling is mediated by IL-17, as demonstrated by the normalization of the majority of hematopoietic parameters following deletion of IL-17Ra, consistent with current understanding. For instance, IL-17 has been shown to stimulate myeloid progenitors and early stage erythroid progenitors. Overexpression or exogenous administration of IL-17 stimulates granulopoiesis and the recruitment of both short- and long-term stem cells [15], relocating core erythropoiesis from bone marrow to spleen [16]. Our data are also consistent with previous studies, including studies of human patients with mutations in STAT1, implying that STAT1-activating cytokines such as IFNs inhibit T_H_17 differentiation [29–42]. Much attention has also been paid to a role for IL-27, a cytokine capable of dramatically inhibiting T_H_17 cell commitment in a STAT1-dependent manner [43]. However, our data argue for a dominant role for IFNs over IL-27, since the proinflammatory phenotype due to STAT1 loss, including enhanced production of IL-17, was mimicked by the combined absence of STAT2 and IFNGR, thereby inactivating IFN-I/III and blocking IFN*γ* responses, but not impairing the action of IL-27, which does not rely on STAT2.

T_H_17 cells differentiate as a spectrum of phenotypically similar cells from homeostatic protective functions to inflammatory pathogenic cells [30, 44, 45]. Consistent with our data, it has been suggested that STAT1 acts at an early stage of T_H_17 differentiation to impair pathogenesis [46], although its molecular targets and the mechanisms regulating T_H_17 pathogenic potential remain incompletely understood. Interestingly, analysis of a published T_H_17 single-cell expression dataset [47] indicated that IFN-stimulated genes are largely restricted to non-pathogenic cells. A possible explanation for altered gene expression in mutant cells is that the absence of STAT1 strengthens signaling by other STATs, resulting in biased T_H_ differentiation. Direct antagonistic actions between STAT1 and STAT3 [48, 49] and between STAT1 and STAT4 [50] have been described. Over-activation of STAT3 or STAT4 due to absence of normal competition from STAT1 could partially underlie the increases in IL-17 and IFN-II observed in KO T cells. However, the fact that loss of STAT2 recapitulates most if not all of the symptoms of a STAT1 KO mouse would argue that such a competition mechanism is at most a minor contributor to the inflammatory process.

Loss of peripheral T_reg_ cells was a striking phenotype of STAT1 KO mice. This loss did not appear to result from impaired T_reg_ differentiation, since no defects were observed *in vitro* or in the thymus. Peripheral T_reg_ loss could reflect a defect in homing to or survival in peripheral lymphoid organs and/or tissues. Decreased homing could be related to the drastic reduction of CD62L, a crucial lymphoid homing molecule [51], observed on STAT1 deficient T cells (Fig. 1). However, it is quite likely that other cell type or the microbiota also regulate the T_reg_ pool [52, 53].

Impaired regulation of T cells in absence of IFN signaling contributes to the propensity for inflammation in STAT1 KO mice, but the phenotype is dependent on a trigger derived from a dysbiotic microbiome, suggesting a major role for tonic IFN in sculpting commensal diversity [12, 54]. How IFN can regulate the microbial population in the gut is still under investigation; however, we hypothesize that this could be achieved in a number of ways. (i) IFN could control gut microbiota through its multiple effects on the immune system, such as by modulating macrophage-mediated bacterial clearance [55]. (ii) TLR recognition of commensal derived products has been shown to be essential for intestinal epithelial homeostasis [56]; since a number of TLR and TLR adapters are IFN inducible [57], it is likely that tonic IFN favors this process. (iii) IFN promotes intestinal epithelial barrier function by enhancing epithelial cell differentiation [54], potentially protecting against bacterial invasion. (iv) IFN has been suggested to have direct antimicrobial properties [58], although these effects may be only indirectly STAT1 dependent, for instance, through a positive feedback loop whereby STAT1-dependent signals amplify IFN expression [59]. (v) IFN alters the expression of hundreds of target genes, possibly rewiring cell metabolism and altering metabolite output [60], thereby modifying the gut microbiota microenvironment. Interestingly, microbiota dysbiosis has also been associated with human diseases where the IFN system is perturbed, such as systemic lupus erythematosus [61].

We found that the gut microbiota of STAT1 deficient mice is depleted of *Bacteroidales S24-7*, which has been described as a beneficial commensal in a mouse model of preclinical inflammatory arthritis [62]. This depletion was accompanied by a concomitant expansion of other families of bacteria, among them *Bacteroidales Prevotellaceae*, in particular species closely related to *P. heparinolytica*. These same changes in the composition of the microbiome were noted in GR STAT2 DKO mice, which phenocopied the inflammatory disease characteristics of STAT1 KO mice. However, these changes were not observed in mouse strains with only partial impairment of IFN signaling that did not phenocopy STAT1 KO. Of note, a number of *Prevotella sp.*, including *P. copri, P. bivia, P. heparinolytica, P. nigrescens* and *P. intermedia*, has been found to drive inflammation through induction of IL-17. *Prevotella sp*. have also been tied to a number of human chronic inflammatory diseases, such as new-onset rheumatoid arthritis (NORA), periodontitis, asthma, and bacterial vaginosis [63–66]. Nevertheless, we cannot rule out a possible contribution of additional bacterial species or of the commensal virome in the observed phenotype [67].

Our data raise the question of whether the altered microbiome of STAT1 KO mice is sufficient to trigger inflammatory disease. Since the microbiome of mice is readily transferred by cohabitation due to coprophagy [68, 69], we have cohoused STAT1 KO mice with WT mice but observed no alteration in the phenotypes of either strain. In addition, we maintained the STAT1 KO colony by interbreeding heterozygous mice, resulting in litters of mixed genotypes, an effective method of allowing redistribution of microbiome composition [70]. However, only homozygous KO offspring in these mixed cohorts developed inflammatory disease. We speculate that competent IFN signaling in WT or heterozygous mice is sufficient to prevent dysbiosis.

The susceptibility of STAT1 KO mice to development of IBD is consistent with the observation that human patients carrying a hypomorphic allele of STAT1 develop symptoms of colitis [7, 8] and that many genes associated with IBD susceptibility are related to the IFN pathway [71]. IFN has been used as a therapy for IBD, but conflicting results have been reported concerning its efficacy in both ulcerative colitis and Crohn’s disease, ranging from disease exacerbation to spectacular remissions [72, 73]. Further studies leading to a better understanding of the molecular mechanisms by which IFN controls gut microbiota to avert inflammatory pathologies will be invaluable for defining the set of patients and the timing where IFN therapy could be beneficial.

The role of IFNs in curbing over-exuberant IL-17 driven inflammation by modulating the microbiota could be of clinical importance in a number of other diseases, including cancer [74]. For instance, STAT1 functions as a tumor suppressor through multiple mechanisms [75]; prevention of inflammation through modulation of the microbiota may provide another facet of its barrier to tumorigenesis. In sum, a delicate balance involving critical homeostatic IFN signaling aids in maintaining a healthy microbiome and taming systemic inflammation. Recent reports have emphasized the importance of gut microbiota on other organs, such as the central nervous system and the lungs [76, 77], suggesting that IFN may have a more far-reaching influence on overall human health and homeostasis through microbiome modulation than currently envisioned.

## Materials and Methods

### Mice

The generation of STAT1-/-, IFNAR-/-, IFNGR-/-, STAT2-/- and IL-17Ra-/- mice has been previously described [78–82]. Mice were interbred to generate doubly or triply gene deficient mice, as indicated, and genotypes were monitored by gene-specific PCR assays. All strains with the exception of STAT1-/- were maintained as homozygous mutant animals. The STAT1 colony was maintained by harem interbreeding of STAT1+/- heterozygous mice, and WT and STAT1-/- mice produced by this cross were compared. Except where indicated in specific experiments that comparisons involved littermates, WT and STAT1-/- offspring were pooled from several litters for experimental procedures. Mice were monitored for disease by inspection of coat condition, posture, enlarged peritoneal cavity indicative of splenomegaly, and peripheral blood counts using a Hemavet 950FS (Drew Scientific) on blood collected from the submandibular vein. All animals used in these experiments were maintained in a single dedicated room of a specific pathogen-free vivarium at NYU Grossman School of Medicine. All work with experimental animals was in accordance with protocols approved by the NYU Langone Health Institutional Animal Care and Use Committee. When indicated, animals were given an antibiotic cocktail made of 1g/L each of ampicillin (Crystalgen), neomycin sulfate (Enzo), and metronidazole (Fisher) and 0.5g/L of vancomycin hydrochloride (Fisher) in the drinking water, as previously described [56]. Where noted, mice were injected weekly with 150mg/Kg of 5- fluorouracil (5-FU) for survival studies. Where indicated, mice were given 3% dextran sulfate sodium (DSS) (M.W. 36.000-50.000, ICN) in drinking water for 7 d, changing the water every 2 d.

## Scoring of disseminated intestinal bacteria

After sacrifice, liver, mesenteric lymph node and spleen lysates were plated on non-selective blood agar plates after serial dilution. For each organ, growth from the neat extract was scored as 1, growth after a 1:10 or 1:100 dilution was scored as 2 or 3, respectively. Final score for each individual mouse was obtained by adding scores obtained for each organ.

### Histological scoring

Postmortem, the entire colon was removed, from the cecum to the anus, and the colon length was measured as a marker of inflammation. The entire colon was fixed in 10% formalin and paraffin sections were stained with hematoxylin and eosin (H&E). Histological scoring was performed in a blinded fashion by a pathologist, with individual scores for inflammatory cell infiltration severity (1: minimal, 2: mild, 3: moderate) and extent (1: expansion into mucosal tissue, 2: expansion to both mucosal and submucosal tissues, 3: mucosal, submucosal and transmural invasion) as well as epithelial damage (1: minimal hyperplasia, 2-3: mild hyperplasia and 3-4: moderate hyperplasia extending to up to 50% of epithelium) and goblet cell loss (scored 1 to 4 depending on severity) [83].

### Flow Cytometry Analysis

Single-cell suspensions were derived from mechanical disruption of mouse bone marrow and spleen in PBS supplemented with 2% FCS (Sigma). For FACS analysis of spleen, bone marrow and peripheral blood, red blood cells were lysed with Pharm Lyse Lysing buffer according to the manufacturer’s instructions (BD Life Sciences). For progenitor staining, bone marrow and spleen cells were incubated with biotin-conjugated antibodies against committed lineage epitopes for 45 min on ice, washed once and incubated with streptavidin and primary antibodies overnight on ice. For mature cell staining, cells were treated with Fc block for 5 min on ice, washed once and incubated with primary antibodies for 45 min on ice. The antibodies used in this study are listed in Table S1. BD LSRII or BD FACSymphony (BD Life Sciences, San Jose, CA) were used for cell acquisition and data were analyzed using FlowJo software (Treestar, Ashland, OR).

### Cytokine Intracellular Staining

Peripheral blood cells were plated after red blood cell lysis in DMEM 10% serum containing 0.2% cell stimulation cocktail (eBiosciences) for 4 h at 37°C. Next, cells were collected, washed once, treated with Fc block for 5 minutes on ice and stained with surface antibodies for 30 minutes on ice. Then, cells were washed, fixed in 2% PFA for 15 min at room temperature, washed again and permeabilized with 0.5% saponin for 10 min at room temperature. Cells were then treated with Fc block diluted in 0.5% saponin for 5 min and stained with intracellular antibodies diluted in 0.5% saponin for 30 min at room temperature. Then cells were washed and incubated with 0.5% saponin for an additional 15 min and analyzed by flow cytometry. The antibodies used in this study are listed in Table S1. BD LSRII (BD Life Sciences, San Jose, CA) was used for all analyses and data were analyzed using FlowJo software (Treestar, Ashland, OR). Gating strategies are shown in Fig. S1.

### Bone marrow and spleen transplant assays

2 X 10^6^ bone marrow or spleen cells harvested from donor mice of appropriate genotypes were intravenously injected in the retro-orbital sinus of lethally irradiated (2 × 5.7 Gy) CD45.1^+^ 8-wk-old recipients. Donor spleens were depleted of T cells prior injection in recipient mice. Briefly, 1 X 10^8^ cells were incubated with anti-CD8-Biotin and anti-CD4-Biotin for 15 min on ice. After one wash, cells were incubated with streptavidin-conjugated Magna beads (Biolegend) for 15 min on ice. The resulting T cell-depleted samples were injected in irradiated recipients. After 6 wk, 2 × 10^6^ whole bone marrow or spleen cells from primary recipient mice were transplanted into lethally irradiated CD45.1^+^ secondary recipient mice. Irradiated mice were given water supplemented with antibiotics for 4 wk. After 4 mo, secondary recipients were analyzed for lineage distribution.

### Colony Formation Unit Assay (methylcellulose)

Total bone marrow and spleen cells from WT and STAT1 KO mice (6-8 weeks) were plated in methylcellulose medium (Methocult M3434, Stem Cell Technologies). Cells were seeded at 20,000 total bone marrow cells and 100,000 total spleen cells per replicate. Colony forming units were enumerated using a Zeiss Axio Observer microscope and re-plated (2,000 cells/replicate) every 7–10 days.

### Multiplex Cytokine Analysis

Cytokines and chemokines in serum were measured by the ProcartaPlex assay (mouse ProcartaPlex panel 1A, Invitrogen), performed on a Luminex 200 machine (Luminex). A panel of 36 cytokines and chemokines were analyzed: CXCL1 (C-X-C motif chemokine 1), CXCL2, CXCL5, CXCL10, CCL2 (C-C motif ligand 2), CCL3, CCL4, CCL5, CCL7, Eotaxin, IFN-*α* (interferon-*α*), IFN-*γ*, IL-1*α* (interlukin-1α), IL-1*β*, IL-2, IL-3, IL-4, IL-5, IL-6, IL-9, IL-10, IL-12p70, IL-13, IL-15/IL-15R, IL-17A, IL-18, IL-22, IL-23, IL-27, IL-28, IL-31, G-CSF (granulocyte colony-stimulating factor), GM–CSF (granulocyte-macrophage colony-stimulating factor), M-CSF (macrophage colony-stimulating factor), LIF (leukemia inhibitory factor), and TNF-*α* (tumor necrosis factor-*α*), according to the manufacturer’s instructions. Cytokine concentrations were determined from the appropriate standard curves of known concentrations of recombinant mouse cytokines and chemokines to convert fluorescence units to concentrations (pg/mL). Each sample was run in duplicate and the mean of the duplicates was used to calculate the measured concentration.

### Gut Microbiome Analysis

#### Gut Microbiota sampling and DNA isolation

Mice fecal pellets were collected from individual mice in each group and stored immediately at -80°C. All animals were aged-matched females, and each genotype was housed separately prior to fecal collection. DNA from fecal pellets was isolated using the Power Soil Kit*^®^* (Qiagen), according to the manufacturer’s instructions, and was stored at -80°C prior to library preparation.

#### 16S rRNA gene sequence analysis

The 16S ribosomal RNA (16S rRNA) V4- region was amplified by using the primers F515-R806, and the products for each fecal sample library were sequenced on a MiSeq instrument (Illumina), as previously described [84]. DNA sequences were analyzed using QIIME 2 as previously described [85, 86]. Operational Taxonomic Units (OTUs) and Amplicon Sequence Variants (ASVs) were obtained using the Dada2 plug-in and assigned using the GreenGenes database to obtain a taxonomy table.

For bacterial community visualization, R package Phyloseq was used to calculate the α-diversity index. Shannon index, Simpson index and observed ASV abundance were used to estimate the community evenness and richness. Kruskal-Wallis test was used to obtain the overall *p* value of the α-diversity index between groups. Phylogenetic analysis was performed using MegAlign Pro (DNA Star, Madison, WI).

### Murine T cell isolation and culture

CD4^+^CD62L^+^ naïve T cells were isolated using CD4 negative enrichment kits (#19852, Stemcell Technologies, Vancouver, Canada) and CD62L microbeads (#130-049-701, Miltenyi Biotec, San Diego, CA) and confirmed to be >95% pure by flow cytometry. These cells were cultured on 96 well plates pre-coated with anti-CD3 and anti-CD28 (3μg/ml each for T_H_17 cultures, 1μg/ml for T_reg_ cultures). Cells were cultured in DMEM supplemented with neutralizing antibodies against IL-12, IFN-*γ* and IL-4 (clones C17.8, XMG1.2 and 11B11, 10μg/ml each, BioXcell, West Lebanon, NH). T_reg_ and T_H_17 cultures were supplemented with TGF-*β* and IL-6 (Peprotech, Rocky Hill, NJ) as indicated. T_reg_ cultures were fed with equal volume of IL-2-supplemented media (10 ng/ml, Peprotech) at day 2, split 1:2 into IL-2-supplemented media at day and analyzed at day 4. T_H_17 cultures were treated similarly except no IL-2 was added.

Five hours prior to analysis, T_H_17 cultures were re-stimulated with PMA and ionomycin (50 and 500 ng/mL, respectively, Sigma Aldrich, St. Louis, MO) in the presence of Golgistop (BD Life Sciences, San Jose, CA). Cells were typically stained with LIVE/DEAD (Thermo Fisher) and anti-CD4-FITC (RM4-5, Biolegend, San Diego, CA) before being fixed and permeabilized using Foxp3 fixation/permeabilization buffers (eBioscience, San Diego, CA). Intracellular staining with anti-IL-17 and anti- FOXP3 (clones eBio17B7 and FJK-16s, eBioscience) was performed per manufacturer’s instructions. Acquisition was performed on a FacsVerse (BD Life Sciences, San Jose, CA) and analyzed using Flowjo software (Treestar, Ashland, OR).

### Statistical analysis

Differences between experimental groups were analyzed for statistical significance using unpaired two-tailed Student’s *t*-test or Mann-Whitney *U* test, where indicated. Data shown are representative of at least three independent experiments, with specific details shown in each figure legend. A *p* < 0.05 was considered to be statistically significant.

## Acknowledgements

The authors thank L. Hu and E. Maucotel for expert technical assistance and G. David, F. Boccalatte, and J. A. Hall for helpful discussions, gift of reagents and support. The authors gratefully acknowledge assistance from the NYU Histopathology, Flow Cytometry, and Microscopy core laboratories, especially C. Loomis, M. Cammer, and M. Gregory. The authors thank Amgen, Inc., for the kind gift of IL-17Ra KO mice. This work was supported in part by National Institutes of Health grants R01AI28900 to DEL, R01AI133822 to SSW, and P50AR070591 to GJS, and by the Laura and Isaac Perlmutter Comprehensive Cancer Center support grant P30CA016087 from the National Cancer Institute.

## Author contributions

IJM conceived the project, designed, performed and analyzed experiments and wrote the manuscript; LB performed and analyzed experiments and helped finalize the manuscript; DA performed and analyzed experiments for the microbiome study; ZC analyzed and performed the bioinformatic analysis of the microbiome study; GB performed and analyzed the multiparameter flow cytometry data; HSL performed the multiplex cytokine analysis; LC provided help and reagents for stem cell analysis; AA performed experiments for analysis of T cell differentiation; WL analyzed histopathology specimens; PC gave advice on the multiparameter flow cytometry analysis and provided access to instrumentation; BK designed and analyzed T_H_17 and T_reg_ differentiation; GJS helped design the microbiome study and obtained funding; SSW suggested and provided advice for the multiplex cytokine analysis and obtained funding; DEL designed and analyzed experiments, wrote the manuscript, provided project supervision, and obtained funding.

## Supplementary Materials

**Fig S1:**
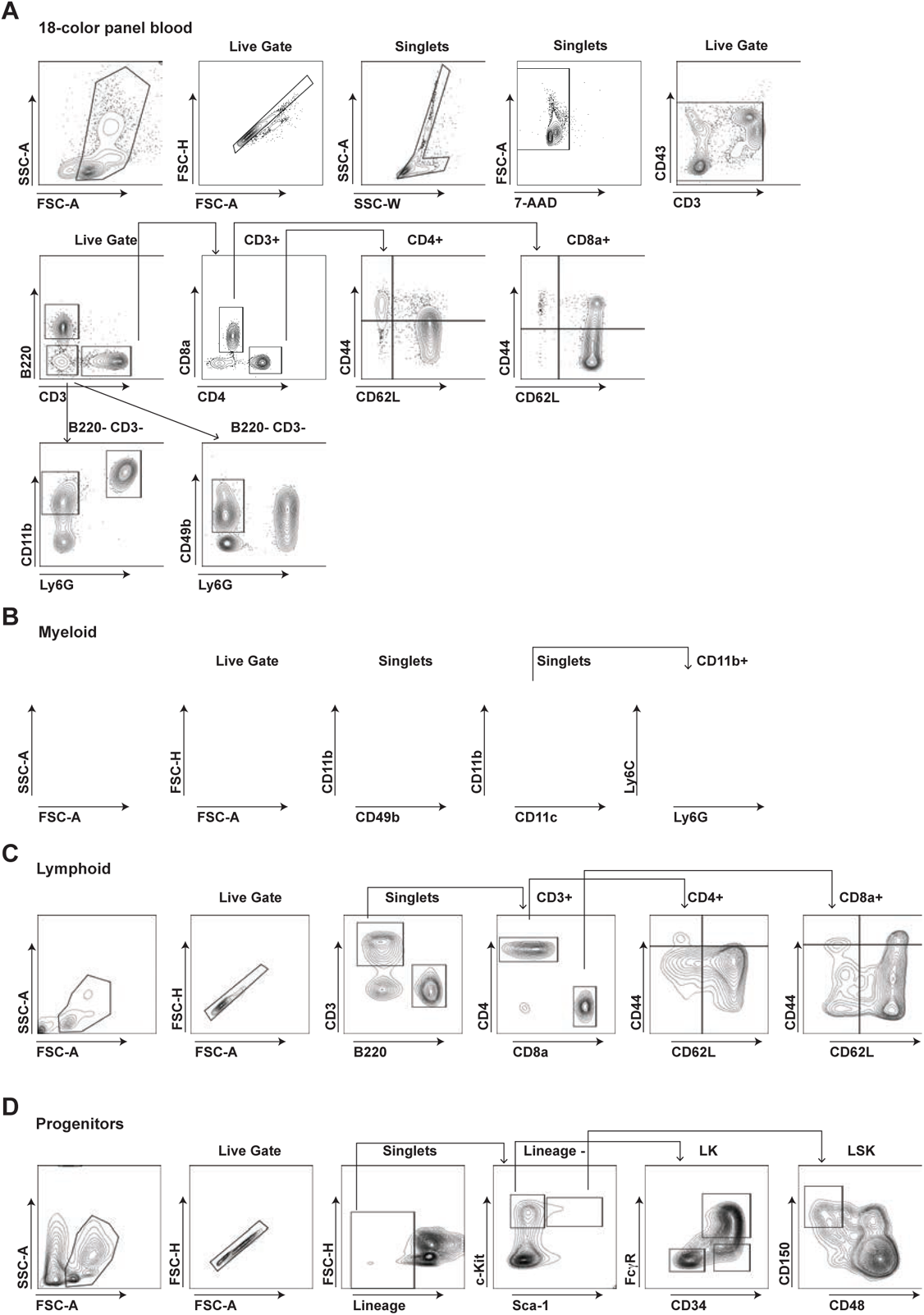
Gating strategies of multi-parameter flow cytometry analysis on blood (A) and for myeloid (B), lymphoid (C) and progenitor subsets (D) from bone marrow and spleen.

**Fig. S2:**
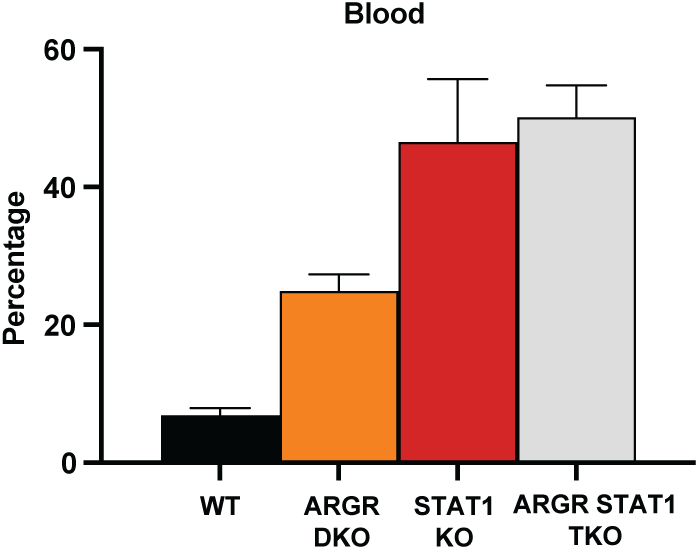
Flow cytometric analysis of myeloid cells in WT, STAT1 KO, ARGR DKO and ARGR STAT1 TKO animals

**Fig. S3:**
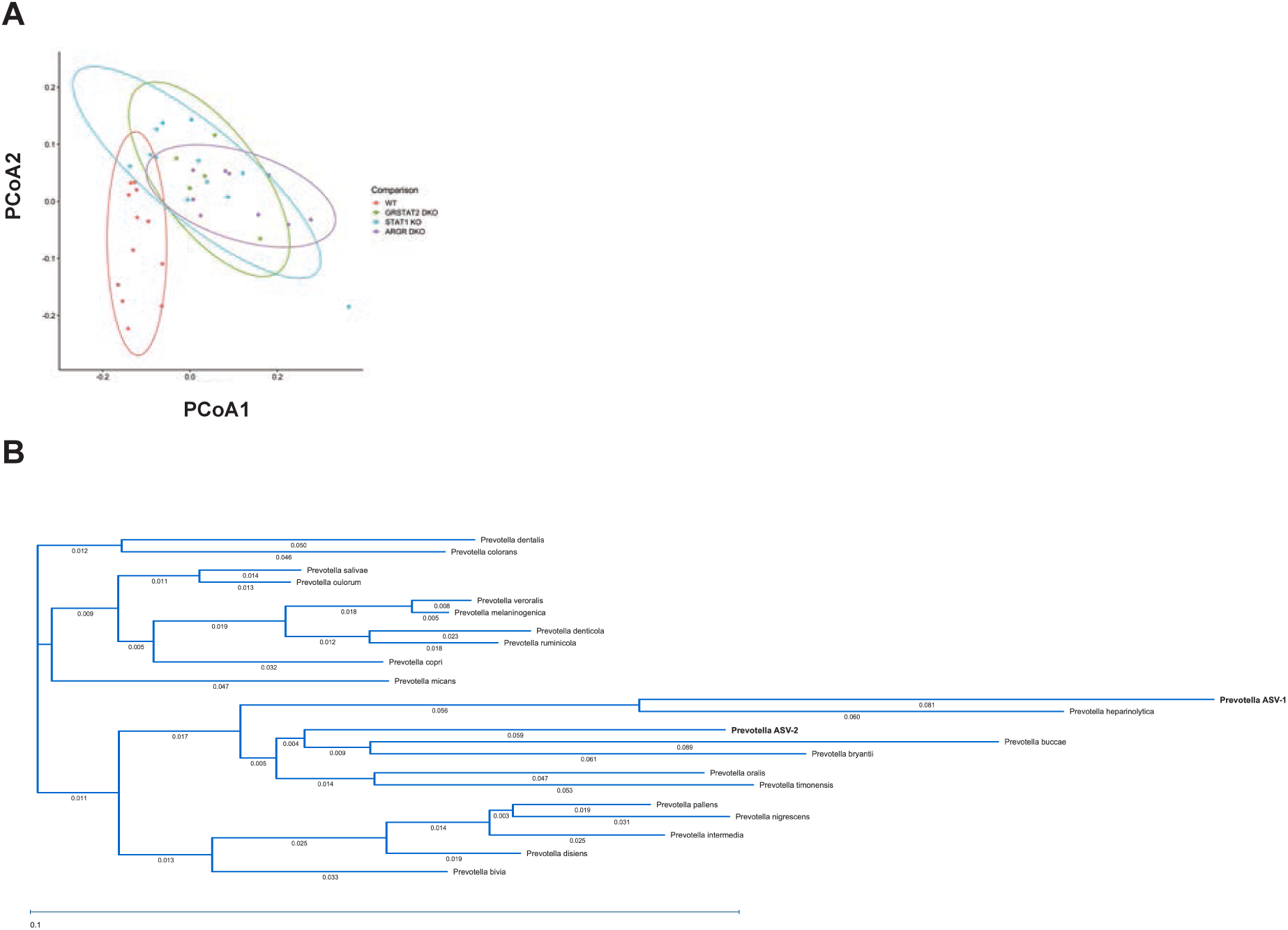
Bioinformatic analysis of beta diversity and phylogeny of *Prevotella* species A- Principal component analysis B- Phylogenetic tree of *Prevotella* species

**Fig. S4:**
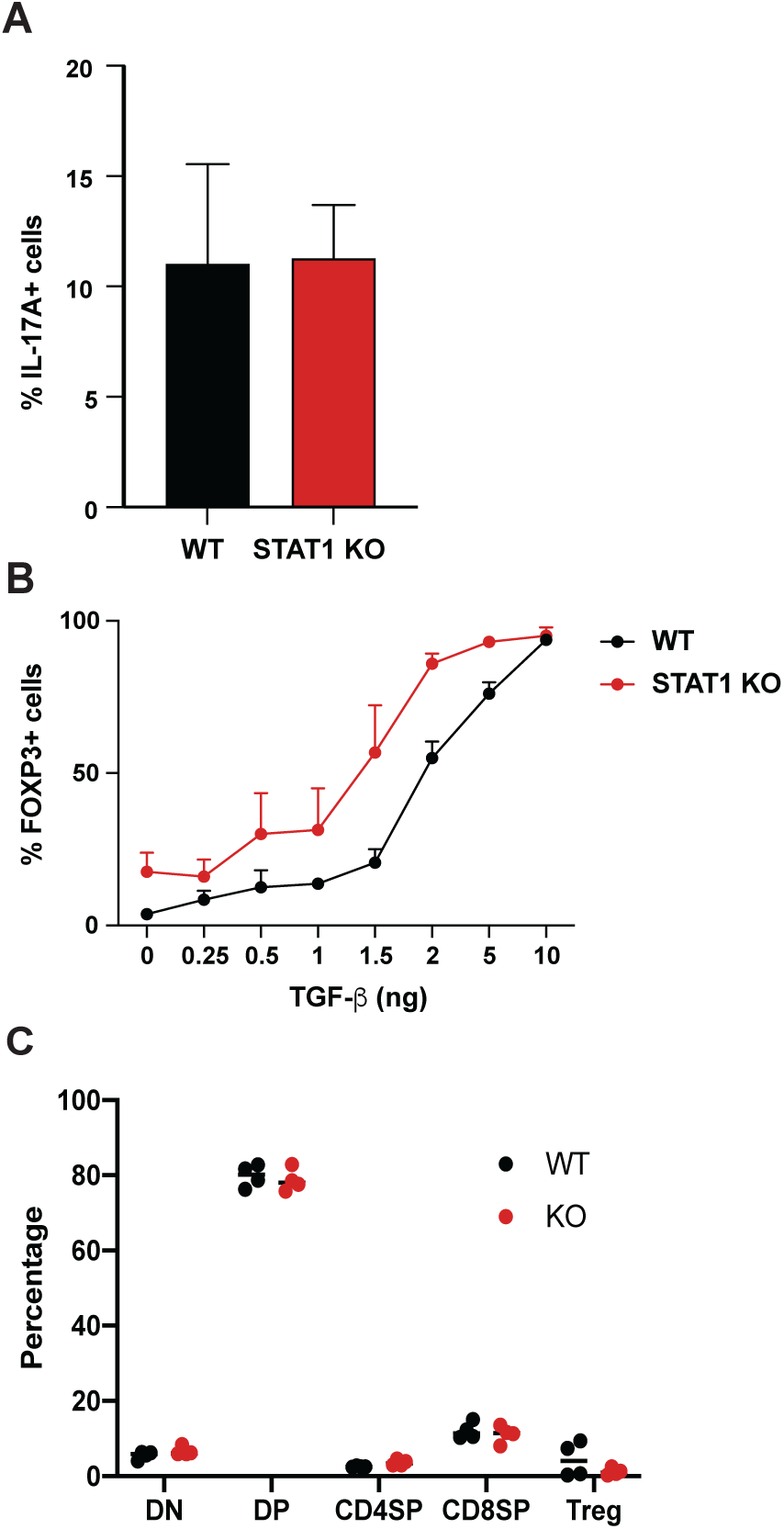
Analysis of STAT1 KO and WT T_H_17 and T_reg_ cells. A- *In vitro* T_H_17 differentiation in response to IL-6 and TGF-β of WT and STAT1 KO T cells (n=3 mice) B- *In vitro* T_reg_ differentiation in response to TGF-β of WT and STAT1 KO T cells (n=3 mice) C- Quantification of thymic T cell populations in WT and STAT1 KO mice (n=4 mice)

**Fig. S5:**
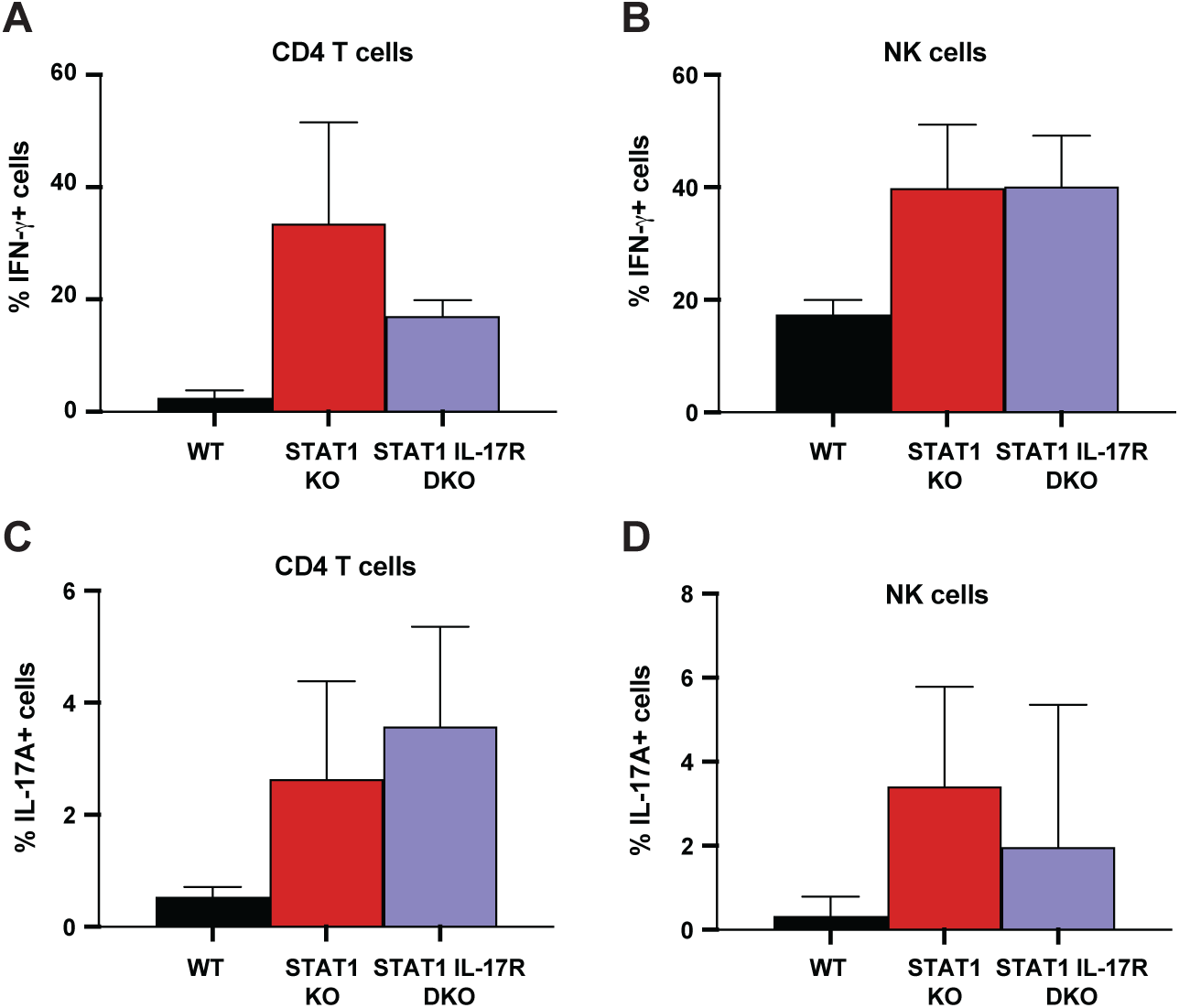
Cytokine production by STAT1 IL-17R DKO lymphocytes Percentage of IFN-γ- (A) or IL-17A-producing (C) CD4+ T cells or NK cells (B and D) in WT, STAT1 KO and IL-17Ra STAT1 DKO mice (n=4 or 5)

**Table S1.**
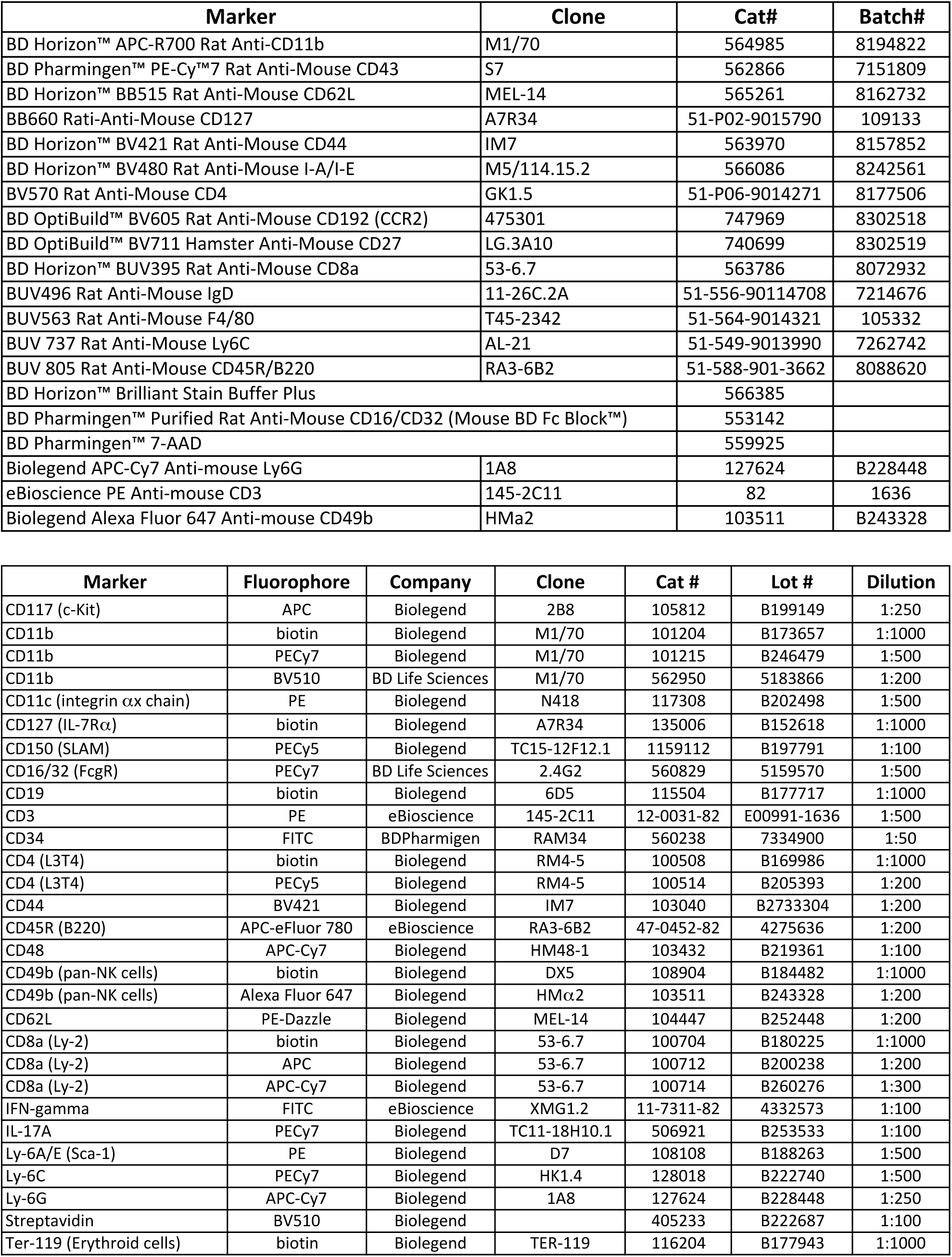
List of antibodies

**Table S2.**
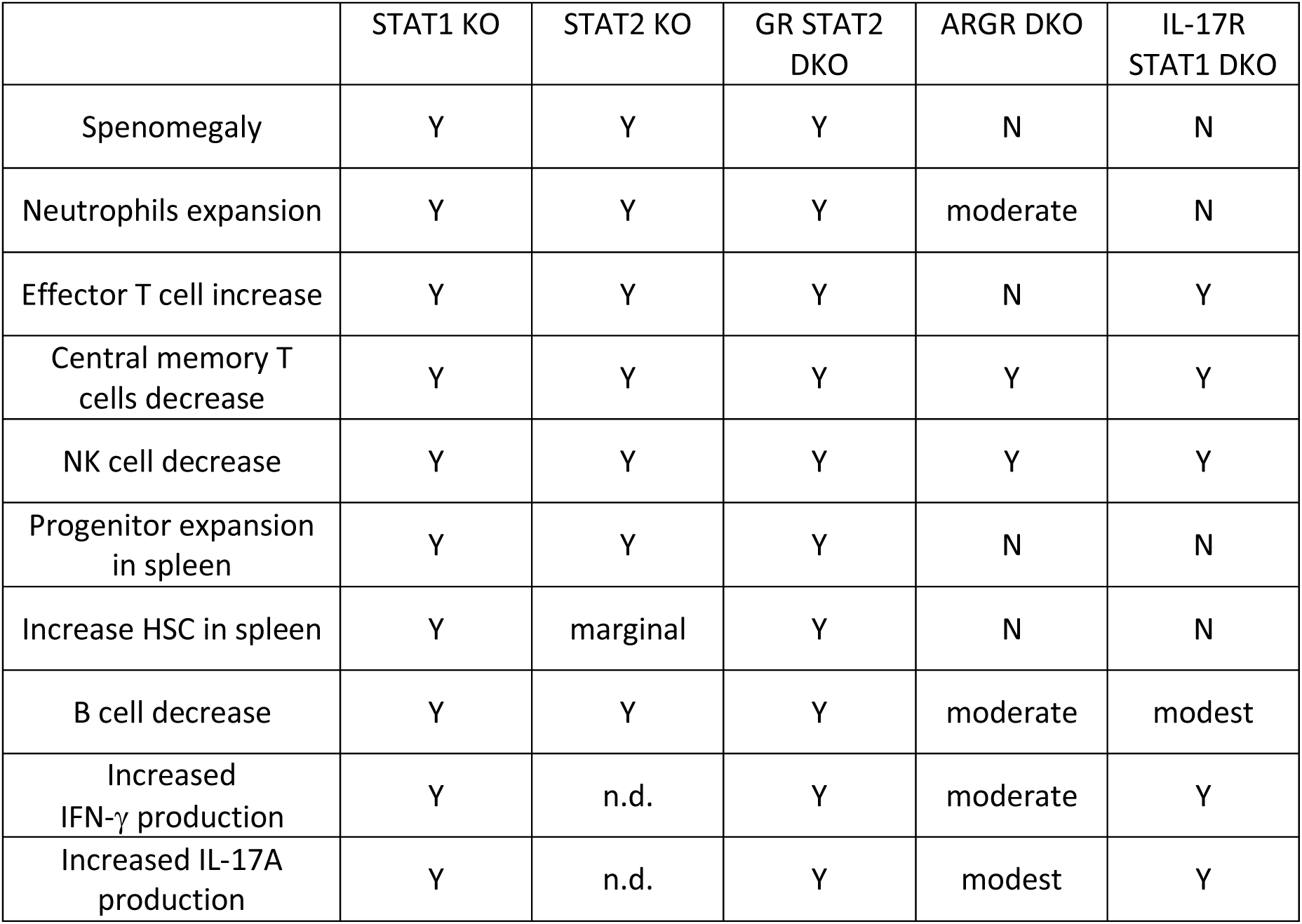
Summary for mutant phenotypes

|  | STAT1 KO | STAT2 KO | GR STAT2 DKO | ARGR DKO | IL-17R STAT1 DKO |
| --- | --- | --- | --- | --- | --- |
| Splenomegaly | Y | Y | Y | N | N |
| Neutrophils expansion | Y | Y | Y | moderate | N |
| Effector T cell increase | Y | Y | Y | N | Y |
| Central memory T cells decrease | Y | Y | Y | Y | Y |
| NK cell decrease | Y | Y | Y | Y | Y |
| Progenitor expansion in spleen | Y | Y | Y | N | N |
| Increase HSC in spleen | Y | marginal | Y | N | N |
| B cell decrease | Y | Y | Y | moderate | modest |
| Increased IFN-γ production | Y | n.d. | Y | moderate | Y |
| Increased IL-17A production | Y | n.d. | Y | modest | Y |

**Data File S1.** Luminex cytokine and chemokine data

Excel file Luminex_STAT1_KO.xls

## References

1. Au-Yeung N, Mandhana R & Horvath CM (2013) Transcriptional regulation by STAT1 and STAT2 in the interferon JAK-STAT pathway. Jak-Stat 2, e23931, doi: 10.4161/jkst.23931.

2. Lazear HM, Schoggins JW & Diamond MS (2019) Shared and Distinct Functions of Type I and Type III Interferons. Immunity 50, 907–923, doi: 10.1016/j.immuni.2019.03.025.

3. Ivashkiv LB (2018) IFNgamma: signalling, epigenetics and roles in immunity, metabolism, disease and cancer immunotherapy. Nat Rev Immunol 18, 545–558, doi: 10.1038/s41577-018-0029-z.

4. Dupuis S, Dargemont C, Fieschi C, Thomassin N, Rosenzweig S, Harris J, Holland SM, Schreiber RD & Casanova JL (2001) Impairment of mycobacterial but not viral immunity by a germline human STAT1 mutation. Science 293, 300–303.

5. Dupuis S, Jouanguy E, Al-Hajjar S, Fieschi C, Al-Mohsen IZ, Al-Jumaah S, Yang K, Chapgier A, Eidenschenk C, Eid P, Al Ghonaium A, Tufenkeji H, Frayha H, Al-Gazlan S, Al-Rayes H, Schreiber RD, Gresser I & Casanova JL (2003) Impaired response to interferon-alpha/beta and lethal viral disease in human STAT1 deficiency. Nat Genet 33, 388–391.

6. Bustamante J, Boisson-Dupuis S, Abel L & Casanova JL (2014) Mendelian susceptibility to mycobacterial disease: genetic, immunological, and clinical features of inborn errors of IFN-gamma immunity. Semin Immunol 26, 454–470, doi: 10.1016/j.smim.2014.09.008.

7. Thoeni C, Hamilton EA, Elkadri A, Murchie R, Fiedler K, Ovadia A, Sharfe N, Ngan B, Cutz E, Nahum A, Muise AM & Roifman CM (2015) The effects of STAT1 dysfunction on the gut. Lymphosign J 3, 19–33, doi: 10.14785/lpsn-2015-0012.

8. Sharfe N, Nahum A, Newell A, Dadi H, Ngan B, Pereira SL, Herbrick JA & Roifman CM (2014) Fatal combined immunodeficiency associated with heterozygous mutation in STAT1. J Allergy Clin Immunol 133, 807–817, doi: 10.1016/j.jaci.2013.09.032.

9. Liu L, Okada S, Kong XF, Kreins AY, Cypowyj S, Abhyankar A, Toubiana J, Itan Y, Audry M, Nitschke P, Masson C, Toth B, Flatot J, Migaud M, Chrabieh M, Kochetkov T, Bolze A, Borghesi A, Toulon A, Hiller J, Eyerich S, Eyerich K, Gulacsy V, Chernyshova L, Chernyshov V, Bondarenko A, Grimaldo RM, Blancas-Galicia L, Beas IM, Roesler J, Magdorf K, Engelhard D, Thumerelle C, Burgel PR, Hoernes M, Drexel B, Seger R, Kusuma T, Jansson AF, Sawalle-Belohradsky J, Belohradsky B, Jouanguy E, Bustamante J, Bue M, Karin N, Wildbaum G, Bodemer C, Lortholary O, Fischer A, Blanche S, Al-Muhsen S, Reichenbach J, Kobayashi M, Rosales FE, Lozano CT, Kilic SS, Oleastro M, Etzioni A, Traidl-Hoffmann C, Renner ED, Abel L, Picard C, Marodi L, Boisson-Dupuis S, Puel A & Casanova JL (2011) Gain-of-function human STAT1 mutations impair IL-17 immunity and underlie chronic mucocutaneous candidiasis. J Exp Med 208, 1635–1648, doi: 10.1084/jem.20110958.

10. Puel A (2020) Human inborn errors of immunity underlying superficial or invasive candidiasis. Hum Genet 139, 1011–1022, doi: 10.1007/s00439-020-02141-7.

11. Broggi MAS, Maillat L, Clement CC, Bordry N, Corthesy P, Auger A, Matter M, Hamelin R, Potin L, Demurtas D, Romano E, Harari A, Speiser DE, Santambrogio L & Swartz MA (2019) Tumor-associated factors are enriched in lymphatic exudate compared to plasma in metastatic melanoma patients. J Exp Med 216, 1091–1107, doi: 10.1084/jem.20181618.

12. Pott J & Stockinger S (2017) Type I and III Interferon in the Gut: Tight Balance between Host Protection and Immunopathology. Front Immunol 8, 258, doi: 10.3389/fimmu.2017.00258.

13. Blazek K, Eames HL, Weiss M, Byrne AJ, Perocheau D, Pease JE, Doyle S, McCann F, Williams RO & Udalova IA (2015) IFN-lambda resolves inflammation via suppression of neutrophil infiltration and IL-1beta production. J Exp Med 212, 845–853, doi: 10.1084/jem.20140995.

14. Krstic A, Mojsilovic S, Jovcic G & Bugarski D (2012) The potential of interleukin- 17 to mediate hematopoietic response. Immunol Res 52, 34–41, doi: 10.1007/s12026-012- 8276-8.

15. Lubberts E, Joosten LA, Oppers B, van den Bersselaar L, Coenen-de Roo CJ, Kolls JK, Schwarzenberger P, van de Loo FA & van den Berg WB (2001) IL-1-independent role of IL-17 in synovial inflammation and joint destruction during collagen-induced arthritis. J Immunol 167, 1004–1013, doi: 10.4049/jimmunol.167.2.1004.

16. Krstic A, Santibanez JF, Okic I, Mojsilovic S, Kocic J, Jovcic G, Milenkovic P & Bugarski D (2010) Combined effect of IL-17 and blockade of nitric oxide biosynthesis on haematopoiesis in mice. Acta Physiol (Oxf) 199, 31–41, doi: 10.1111/j.1748- 1716.2010.02082.x.

17. Belkaid Y & Hand TW (2014) Role of the microbiota in immunity and inflammation. Cell 157, 121–141, doi: 10.1016/j.cell.2014.03.011.

18. Gutierrez-Merino J, Isla B, Combes T, Martinez-Estrada F & Maluquer De Motes C (2020) Beneficial bacteria activate type-I interferon production via the intracellular cytosolic sensors STING and MAVS. Gut Microbes 11, 771–788, doi: 10.1080/19490976.2019.1707015.

19. Castillo-Alvarez F, Perez-Matute P, Oteo JA & Marzo-Sola ME (2018) The influence of interferon beta-1b on gut microbiota composition in patients with multiple sclerosis. Neurologia, doi: 10.1016/j.nrl.2018.04.006.

20. Alteber Z, Sharbi-Yunger A, Pevsner-Fischer M, Blat D, Roitman L, Tzehoval E, Elinav E & Eisenbach L (2018) The anti-inflammatory IFITM genes ameliorate colitis and partially protect from tumorigenesis by changing immunity and microbiota. Immunol Cell Biol 96, 284–297, doi: 10.1111/imcb.12000.

21. Canesso MCC, Lemos L, Neves TC, Marim FM, Castro TBR, Veloso ES, Queiroz CP, Ahn J, Santiago HC, Martins FS, Alves-Silva J, Ferreira E, Cara DC, Vieira AT, Barber GN, Oliveira SC & Faria AMC (2018) The cytosolic sensor STING is required for intestinal homeostasis and control of inflammation. Mucosal Immunol 11, 820–834, doi: 10.1038/mi.2017.88.

22. Morita Y, Iseki A, Okamura S, Suzuki S, Nakauchi H & Ema H (2011) Functional characterization of hematopoietic stem cells in the spleen. Exp Hematol 39, 351–359 e353, doi: 10.1016/j.exphem.2010.12.008.

23. Gimeno R, Lee CK, Schindler C & Levy DE (2005) Stat1 and Stat2 but not Stat3 Arbitrate Contradictory Growth Signals Elicited by IFNa/b in T Lymphocytes. Mol Cell Biol 25, 5456–5465.

24. Alkim C, Alkim H, Koksal AR, Boga S & Sen I (2015) Angiogenesis in Inflammatory Bowel Disease. Int J Inflam 2015, 970890, doi: 10.1155/2015/970890.

25. Calcinotto A, Brevi A, Chesi M, Ferrarese R, Garcia Perez L, Grioni M, Kumar S, Garbitt VM, Sharik ME, Henderson KJ, Tonon G, Tomura M, Miwa Y, Esplugues E, Flavell RA, Huber S, Canducci F, Rajkumar VS, Bergsagel PL & Bellone M (2018) Microbiota-driven interleukin-17-producing cells and eosinophils synergize to accelerate multiple myeloma progression. Nat Commun 9, 4832, doi: 10.1038/s41467-018-07305-8.

26. Yamada A, Arakaki R, Saito M, Tsunematsu T, Kudo Y & Ishimaru N (2016) Role of regulatory T cell in the pathogenesis of inflammatory bowel disease. World J Gastroenterol 22, 2195–2205, doi: 10.3748/wjg.v22.i7.2195.

27. Tan W, Huang W, Zhong Q & Schwarzenberger P (2006) IL-17 receptor knockout mice have enhanced myelotoxicity and impaired hemopoietic recovery following gamma irradiation. J Immunol 176, 6186–6193, doi: 10.4049/jimmunol.176.10.6186.

28. Porpaczy E, Tripolt S, Hoelbl-Kovacic A, Gisslinger B, Bago-Horvath Z, Casanova-Hevia E, Clappier E, Decker T, Fajmann S, Fux DA, Greiner G, Gueltekin S, Heller G, Herkner H, Hoermann G, Kiladjian JJ, Kolbe T, Kornauth C, Krauth MT, Kralovics R, Muellauer L, Mueller M, Prchal-Murphy M, Putz EM, Raffoux E, Schiefer AI, Schmetterer K, Schneckenleithner C, Simonitsch-Klupp I, Skrabs C, Sperr WR, Staber PB, Strobl B, Valent P, Jaeger U, Gisslinger H & Sexl V (2018) Aggressive B-cell lymphomas in patients with myelofibrosis receiving JAK1/2 inhibitor therapy. Blood 132, 694–706, doi: 10.1182/blood-2017-10-810739.

29. Batten M, Li J, Yi S, Kljavin NM, Danilenko DM, Lucas S, Lee J, de Sauvage FJ & Ghilardi N (2006) Interleukin 27 limits autoimmune encephalomyelitis by suppressing the development of interleukin 17-producing T cells. Nat Immunol 7, 929–936, doi: 10.1038/ni1375.

30. Yoshimura T, Takeda A, Hamano S, Miyazaki Y, Kinjyo I, Ishibashi T, Yoshimura A & Yoshida H (2006) Two-sided roles of IL-27: induction of Th1 differentiation on naive CD4+ T cells versus suppression of proinflammatory cytokine production including IL- 23-induced IL-17 on activated CD4+ T cells partially through STAT3-dependent mechanism. J Immunol 177, 5377–5385, doi: 10.4049/jimmunol.177.8.5377.

31. Stumhofer JS, Laurence A, Wilson EH, Huang E, Tato CM, Johnson LM, Villarino AV, Huang Q, Yoshimura A, Sehy D, Saris CJ, O’Shea JJ, Hennighausen L, Ernst M & Hunter CA (2006) Interleukin 27 negatively regulates the development of interleukin 17- producing T helper cells during chronic inflammation of the central nervous system. Nat Immunol 7, 937–945, doi: 10.1038/ni1376.

32. Amadi-Obi A, Yu CR, Liu X, Mahdi RM, Clarke GL, Nussenblatt RB, Gery I, Lee YS & Egwuagu CE (2007) T_H_17 cells contribute to uveitis and scleritis and are expanded by IL-2 and inhibited by IL-27/STAT1. Nat Med 13, 711–718, doi: 10.1038/nm1585.

33. Feng G, Gao W, Strom TB, Oukka M, Francis RS, Wood KJ & Bushell A (2008) Exogenous IFN-gamma ex vivo shapes the alloreactive T-cell repertoire by inhibition of T_H_17 responses and generation of functional Foxp3+ regulatory T cells. Eur J Immunol 38, 2512–2527, doi: 10.1002/eji.200838411.

34. Kimura A, Naka T, Nohara K, Fujii-Kuriyama Y & Kishimoto T (2008) Aryl hydrocarbon receptor regulates Stat1 activation and participates in the development of T_H_17 cells. Proc Natl Acad Sci U S A 105, 9721–9726, doi: 10.1073/pnas.0804231105.

35. Tanaka K, Ichiyama K, Hashimoto M, Yoshida H, Takimoto T, Takaesu G, Torisu T, Hanada T, Yasukawa H, Fukuyama S, Inoue H, Nakanishi Y, Kobayashi T & Yoshimura A (2008) Loss of suppressor of cytokine signaling 1 in helper T cells leads to defective T_H_17 differentiation by enhancing antagonistic effects of IFN-gamma on STAT3 and Smads. J Immunol 180, 3746–3756, doi: 10.4049/jimmunol.180.6.3746.

36. Chen M, Chen G, Nie H, Zhang X, Niu X, Zang YC, Skinner SM, Zhang JZ, Killian JM & Hong J (2009) Regulatory effects of IFN-beta on production of osteopontin and IL- 17 by CD4+ T Cells in MS. Eur J Immunol 39, 2525–2536, doi: 10.1002/eji.200838879.

37. Ramgolam VS, Sha Y, Jin J, Zhang X & Markovic-Plese S (2009) IFN-beta inhibits human T_H_17 cell differentiation. J Immunol 183, 5418–5427, doi: 10.4049/jimmunol.0803227.

38. Diveu C, McGeachy MJ, Boniface K, Stumhofer JS, Sathe M, Joyce-Shaikh B, Chen Y, Tato CM, McClanahan TK, de Waal Malefyt R, Hunter CA, Cua DJ & Kastelein RA (2009) IL-27 blocks RORc expression to inhibit lineage commitment of T_H_17 cells. J Immunol 182, 5748–5756, doi: 10.4049/jimmunol.0801162.

39. El-behi M, Ciric B, Yu S, Zhang GX, Fitzgerald DC & Rostami A (2009) Differential effect of IL-27 on developing versus committed T_H_17 cells. J Immunol 183, 4957–4967, doi: 10.4049/jimmunol.0900735.

40. Villarino AV, Gallo E & Abbas AK (2010) STAT1-activating cytokines limit T_H_17 responses through both T-bet-dependent and -independent mechanisms. J Immunol 185, 6461–6471, doi: 10.4049/jimmunol.1001343.

41. Liu H & Rohowsky-Kochan C (2011) Interleukin-27-mediated suppression of human T_H_17 cells is associated with activation of STAT1 and suppressor of cytokine signaling protein 1. J Interferon Cytokine Res 31, 459–469, doi: 10.1089/jir.2010.0115.

42. Okada S, Asano T, Moriya K, Boisson-Dupuis S, Kobayashi M, Casanova JL & Puel A (2020) Human STAT1 Gain-of-Function Heterozygous Mutations: Chronic Mucocutaneous Candidiasis and Type I Interferonopathy. J Clin Immunol 40, 1065–1081, doi: 10.1007/s10875-020-00847-x.

43. Wang Q & Liu J (2016) Regulation and Immune Function of IL-27. Adv Exp Med Biol 941, 191–211, doi: 10.1007/978-94-024-0921-5_9.

44. Gaublomme JT, Yosef N, Lee Y, Gertner RS, Yang LV, Wu C, Pandolfi PP, Mak T, Satija R, Shalek AK, Kuchroo VK, Park H & Regev A (2015) Single-Cell Genomics Unveils Critical Regulators of T_H_17 Cell Pathogenicity. Cell 163, 1400–1412, doi: 10.1016/j.cell.2015.11.009.

45. Ciofani M, Madar A, Galan C, Sellars M, Mace K, Pauli F, Agarwal A, Huang W, Parkhurst CN, Muratet M, Newberry KM, Meadows S, Greenfield A, Yang Y, Jain P, Kirigin FK, Birchmeier C, Wagner EF, Murphy KM, Myers RM, Bonneau R & Littman DR (2012) A validated regulatory network for T_H_17 cell specification. Cell 151, 289–303, doi: 10.1016/j.cell.2012.09.016.

46. Yosef N, Shalek AK, Gaublomme JT, Jin H, Lee Y, Awasthi A, Wu C, Karwacz K, Xiao S, Jorgolli M, Gennert D, Satija R, Shakya A, Lu DY, Trombetta JJ, Pillai MR, Ratcliffe PJ, Coleman ML, Bix M, Tantin D, Park H, Kuchroo VK & Regev A (2013) Dynamic regulatory network controlling T_H_17 cell differentiation. Nature 496, 461–468, doi: 10.1038/nature11981.

47. Karmaus PWF, Chen X, Lim SA, Herrada AA, Nguyen T-LM, Xu B, Dhungana Y, Rankin S, Chen W, Rosencrance C, Yang K, Fan Y, Cheng Y, Easton J, Neale G, Vogel P & Chi H (2019) Metabolic heterogeneity underlies reciprocal fates of T_H_17 cell stemness and plasticity. Nature 565, 101–105, doi: 10.1038/s41586-018-0806-7 PMID - 30568299.

48. Regis G, Pensa S, Boselli D, Novelli F & Poli V (2008) Ups and downs: the STAT1:STAT3 seesaw of Interferon and gp130 receptor signalling. Semin Cell Dev Biol 19, 351–359, doi: 10.1016/j.semcdb.2008.06.004.

49. Wan CK, Andraski AB, Spolski R, Li P, Kazemian M, Oh J, Samsel L, Swanson PA, 2nd, McGavern DB, Sampaio EP, Freeman AF, Milner JD, Holland SM & Leonard WJ (2015) Opposing roles of STAT1 and STAT3 in IL-21 function in CD4+ T cells. Proc Natl Acad Sci U S A 112, 9394–9399, doi: 10.1073/pnas.1511711112.

50. Nguyen KB, Watford WT, Salomon R, Hofmann SR, Pien GC, Morinobu A, Gadina M, O’Shea JJ & Biron CA (2002) Critical role for STAT4 activation by type 1 interferons in the interferon-gamma response to viral infection. Science 297, 2063–2066, doi: 10.1126/science.1074900.

51. Wirth TC, Badovinac VP, Zhao L, Dailey MO & Harty JT (2009) Differentiation of central memory CD8 T cells is independent of CD62L-mediated trafficking to lymph nodes. J Immunol 182, 6195–6206, doi: 10.4049/jimmunol.0803315.

52. Atarashi K, Tanoue T, Shima T, Imaoka A, Kuwahara T, Momose Y, Cheng G, Yamasaki S, Saito T, Ohba Y, Taniguchi T, Takeda K, Hori S, Ivanov, II, Umesaki Y, Itoh K & Honda K (2011) Induction of colonic regulatory T cells by indigenous Clostridium species. Science 331, 337–341, doi: 10.1126/science.1198469.

53. Furusawa Y, Obata Y, Fukuda S, Endo TA, Nakato G, Takahashi D, Nakanishi Y, Uetake C, Kato K, Kato T, Takahashi M, Fukuda NN, Murakami S, Miyauchi E, Hino S, Atarashi K, Onawa S, Fujimura Y, Lockett T, Clarke JM, Topping DL, Tomita M, Hori S, Ohara O, Morita T, Koseki H, Kikuchi J, Honda K, Hase K & Ohno H (2013) Commensal microbe-derived butyrate induces the differentiation of colonic regulatory T cells. Nature 504, 446–450, doi: 10.1038/nature12721.

54. Kotredes KP, Thomas B & Gamero AM (2017) The Protective Role of Type I Interferons in the Gastrointestinal Tract. Front Immunol 8, 410, doi: 10.3389/fimmu.2017.00410.

55. Kumaran Satyanarayanan S, El Kebir D, Soboh S, Butenko S, Sekheri M, Saadi J, Peled N, Assi S, Othman A, Schif-Zuck S, Feuermann Y, Barkan D, Sher N, Filep JG & Ariel A (2019) IFN-beta is a macrophage-derived effector cytokine facilitating the resolution of bacterial inflammation. Nat Commun 10, 3471, doi: 10.1038/s41467-019- 10903-9.

56. Rakoff-Nahoum S, Paglino J, Eslami-Varzaneh F, Edberg S & Medzhitov R (2004) Recognition of commensal microflora by toll-like receptors is required for intestinal homeostasis. Cell 118, 229–241, doi: 10.1016/j.cell.2004.07.002.

57. Siren J, Pirhonen J, Julkunen I & Matikainen S (2005) IFN-alpha regulates TLR-dependent gene expression of IFN-alpha, IFN-beta, IL-28, and IL-29. J Immunol 174, 1932–1937, doi: 10.4049/jimmunol.174.4.1932.

58. Kaplan A, Lee MW, Wolf AJ, Limon JJ, Becker CA, Ding M, Murali R, Lee EY, Liu GY, Wong GCL & Underhill DM (2017) Direct Antimicrobial Activity of IFN-beta. J Immunol 198, 4036–4045, doi: 10.4049/jimmunol.1601226.

59. Marie I, Durbin JE & Levy DE (1998) Differential viral induction of distinct interferon-alpha genes by positive feedback through interferon regulatory factor-7. EMBO J 17, 6660–6669.

60. Fritsch SD & Weichhart T (2016) Effects of Interferons and Viruses on Metabolism. Front Immunol 7, 630, doi: 10.3389/fimmu.2016.00630.

61. Silverman GJ, Azzouz DF & Alekseyenko AV (2019) Systemic Lupus Erythematosus and dysbiosis in the microbiome: cause or effect or both? Curr Opin Immunol 61, 80–85, doi: 10.1016/j.coi.2019.08.007.

62. Rogier R, Evans-Marin H, Manasson J, van der Kraan PM, Walgreen B, Helsen MM, van den Bersselaar LA, van de Loo FA, van Lent PL, Abramson SB, van den Berg WB, Koenders MI, Scher JU & Abdollahi-Roodsaz S (2017) Alteration of the intestinal microbiome characterizes preclinical inflammatory arthritis in mice and its modulation attenuates established arthritis. Sci Rep 7, 15613, doi: 10.1038/s41598-017-15802-x.

63. Scher JU, Sczesnak A, Longman RS, Segata N, Ubeda C, Bielski C, Rostron T, Cerundolo V, Pamer EG, Abramson SB, Huttenhower C & Littman DR (2013) Expansion of intestinal Prevotella copri correlates with enhanced susceptibility to arthritis. Elife 2, e01202, doi: 10.7554/eLife.01202.

64. Larsen JM (2017) The immune response to Prevotella bacteria in chronic inflammatory disease. Immunology 151, 363–374, doi: 10.1111/imm.12760.

65. Si J, You HJ, Yu J, Sung J & Ko G (2017) Prevotella as a Hub for Vaginal Microbiota under the Influence of Host Genetics and Their Association with Obesity. Cell Host Microbe 21, 97–105, doi: 10.1016/j.chom.2016.11.010.

66. Lopes MP, Cruz AA, Xavier MT, Stocker A, Carvalho-Filho P, Miranda PM, Meyer RJ, Soledade KR, Gomes-Filho IS & Trindade SC (2020) Prevotella intermedia and periodontitis are associated with severe asthma. J Periodontol 91, 46–54, doi: 10.1002/JPER.19-0065.

67. Neil JA & Cadwell K (2018) The Intestinal Virome and Immunity. J Immunol 201, 1615–1624, doi: 10.4049/jimmunol.1800631.

68. Soave O & Brand CD (1991) Coprophagy in animals: a review. Cornell Vet 81, 357–364.

69. Caruso R, Ono M, Bunker ME, Nunez G & Inohara N (2019) Dynamic and Asymmetric Changes of the Microbial Communities after Cohousing in Laboratory Mice. Cell Rep 27, 3401–3412 e3403, doi: 10.1016/j.celrep.2019.05.042.

70. Robertson SJ, Lemire P, Maughan H, Goethel A, Turpin W, Bedrani L, Guttman DS, Croitoru K, Girardin SE & Philpott DJ (2019) Comparison of Co-housing and Littermate Methods for Microbiota Standardization in Mouse Models. Cell Rep 27, 1910–1919 e1912, doi: 10.1016/j.celrep.2019.04.023.

71. Jostins L, Ripke S, Weersma RK, Duerr RH, McGovern DP, Hui KY, Lee JC, Schumm LP, Sharma Y, Anderson CA, Essers J, Mitrovic M, Ning K, Cleynen I, Theatre E, Spain SL, Raychaudhuri S, Goyette P, Wei Z, Abraham C, Achkar JP, Ahmad T, Amininejad L, Ananthakrishnan AN, Andersen V, Andrews JM, Baidoo L, Balschun T, Bampton PA, Bitton A, Boucher G, Brand S, Buning C, Cohain A, Cichon S, D’Amato M, De Jong D, Devaney KL, Dubinsky M, Edwards C, Ellinghaus D, Ferguson LR, Franchimont D, Fransen K, Gearry R, Georges M, Gieger C, Glas J, Haritunians T, Hart A, Hawkey C, Hedl M, Hu X, Karlsen TH, Kupcinskas L, Kugathasan S, Latiano A, Laukens D, Lawrance IC, Lees CW, Louis E, Mahy G, Mansfield J, Morgan AR, Mowat C, Newman W, Palmieri O, Ponsioen CY, Potocnik U, Prescott NJ, Regueiro M, Rotter JI, Russell RK, Sanderson JD, Sans M, Satsangi J, Schreiber S, Simms LA, Sventoraityte J, Targan SR, Taylor KD, Tremelling M, Verspaget HW, De Vos M, Wijmenga C, Wilson DC, Winkelmann J, Xavier RJ, Zeissig S, Zhang B, Zhang CK, Zhao H, International IBDGC, Silverberg MS, Annese V, Hakonarson H, Brant SR, Radford-Smith G, Mathew CG, Rioux JD, Schadt EE, Daly MJ, Franke A, Parkes M, Vermeire S, Barrett JC & Cho JH (2012) Host-microbe interactions have shaped the genetic architecture of inflammatory bowel disease. Nature 491, 119–124, doi: 10.1038/nature11582.

72. Nikolaus S, Rutgeerts P, Fedorak R, Steinhart AH, Wild GE, Theuer D, Mohrle J & Schreiber S (2003) Interferon beta-1a in ulcerative colitis: a placebo controlled, randomised, dose escalating study. Gut 52, 1286–1290, doi: 10.1136/gut.52.9.1286.

73. Ferre Aracil C, Rodriguez de Santiago E, Garcia Garcia de Paredes A, Aguilera Castro L, Soria Rivas A & Lopez SanRoman A (2016) [Complete remission of Crohn’s disease after high-dose alpha-interferon treatment for malignant melanoma]. Gastroenterol Hepatol 39, 397–400, doi: 10.1016/j.gastrohep.2015.05.004.

74. Zhao J, Chen X, Herjan T & Li X (2020) The role of interleukin-17 in tumor development and progression. J Exp Med 217, doi: 10.1084/jem.20190297.

75. Meissl K, Macho-Maschler S, Muller M & Strobl B (2017) The good and the bad faces of STAT1 in solid tumours. Cytokine 89, 12–20, doi: 10.1016/j.cyto.2015.11.011.

76. Blacher E, Bashiardes S, Shapiro H, Rothschild D, Mor U, Dori-Bachash M, Kleimeyer C, Moresi C, Harnik Y, Zur M, Zabari M, Brik RB, Kviatcovsky D, Zmora N, Cohen Y, Bar N, Levi I, Amar N, Mehlman T, Brandis A, Biton I, Kuperman Y, Tsoory M, Alfahel L, Harmelin A, Schwartz M, Israelson A, Arike L, Johansson MEV, Hansson GC, Gotkine M, Segal E & Elinav E (2019) Potential roles of gut microbiome and metabolites in modulating ALS in mice. Nature 572, 474–480, doi: 10.1038/s41586-019-1443-5.

77. Budden KF, Gellatly SL, Wood DL, Cooper MA, Morrison M, Hugenholtz P & Hansbro PM (2017) Emerging pathogenic links between microbiota and the gut-lung axis. Nat Rev Microbiol 15, 55–63, doi: 10.1038/nrmicro.2016.142.

78. Durbin JE, Hackenmiller R, Simon MC & Levy DE (1996) Targeted disruption of the mouse Stat1 gene results in compromised innate immunity to viral disease. Cell 84, 443–450.

79. Müller U, Steinhoff U, Reis LFL, Hemmi S, Pavlovic J, Zinkernagel RM & Aguet M (1994) Functional role of type I and type II interferons in antiviral defense. Science 264, 1918–1921.

80. Huang S, Hendriks W, Althage A, Hemmi S, Bluethmann H, Kamijo R, Vilcek J, Zinkernagel RM & Aguet M (1993) Immune response in mice that lack the interferon-gamma receptor. Science 259, 1742–1745.

81. Park C, Li S, Cha E & Schindler C (2000) Immune response in Stat2 knockout mice. Immunity 13, 795–804.

82. Ye P, Rodriguez FH, Kanaly S, Stocking KL, Schurr J, Schwarzenberger P, Oliver P, Huang W, Zhang P, Zhang J, Shellito JE, Bagby GJ, Nelson S, Charrier K, Peschon JJ & Kolls JK (2001) Requirement of interleukin 17 receptor signaling for lung CXC chemokine and granulocyte colony-stimulating factor expression, neutrophil recruitment, and host defense. J Exp Med 194, 519–527, doi: 10.1084/jem.194.4.519.

83. Erben U, Loddenkemper C, Doerfel K, Spieckermann S, Haller D, Heimesaat MM, Zeitz M, Siegmund B & Kuhl AA (2014) A guide to histomorphological evaluation of intestinal inflammation in mouse models. Int J Clin Exp Pathol 7, 4557–4576.

84. Azzouz D, Omarbekova A, Heguy A, Schwudke D, Gisch N, Rovin BH, Caricchio R, Buyon JP, Alekseyenko AV & Silverman GJ (2019) Lupus nephritis is linked to disease-activity associated expansions and immunity to a gut commensal. Ann Rheum Dis 78, 947–956, doi: 10.1136/annrheumdis-2018-214856.

85. Caporaso JG, Kuczynski J, Stombaugh J, Bittinger K, Bushman FD, Costello EK, Fierer N, Pena AG, Goodrich JK, Gordon JI, Huttley GA, Kelley ST, Knights D, Koenig JE, Ley RE, Lozupone CA, McDonald D, Muegge BD, Pirrung M, Reeder J, Sevinsky JR, Turnbaugh PJ, Walters WA, Widmann J, Yatsunenko T, Zaneveld J & Knight R (2010) QIIME allows analysis of high-throughput community sequencing data. Nat Methods 7, 335–336, doi: 10.1038/nmeth.f.303.

86. Bolyen E, Rideout JR, Dillon MR, Bokulich NA, Abnet CC, Al-Ghalith GA, Alexander H, Alm EJ, Arumugam M, Asnicar F, Bai Y, Bisanz JE, Bittinger K, Brejnrod A, Brislawn CJ, Brown CT, Callahan BJ, Caraballo-Rodriguez AM, Chase J, Cope EK, Da Silva R, Diener C, Dorrestein PC, Douglas GM, Durall DM, Duvallet C, Edwardson CF, Ernst M, Estaki M, Fouquier J, Gauglitz JM, Gibbons SM, Gibson DL, Gonzalez A, Gorlick K, Guo J, Hillmann B, Holmes S, Holste H, Huttenhower C, Huttley GA, Janssen S, Jarmusch AK, Jiang L, Kaehler BD, Kang KB, Keefe CR, Keim P, Kelley ST, Knights D, Koester I, Kosciolek T, Kreps J, Langille MGI, Lee J, Ley R, Liu YX, Loftfield E, Lozupone C, Maher M, Marotz C, Martin BD, McDonald D, McIver LJ, Melnik AV, Metcalf JL, Morgan SC, Morton JT, Naimey AT, Navas-Molina JA, Nothias LF, Orchanian SB, Pearson T, Peoples SL, Petras D, Preuss ML, Pruesse E, Rasmussen LB, Rivers A, Robeson MS, 2nd, Rosenthal P, Segata N, Shaffer M, Shiffer A, Sinha R, Song SJ, Spear JR, Swafford AD, Thompson LR, Torres PJ, Trinh P, Tripathi A, Turnbaugh PJ, Ul-Hasan S, van der Hooft JJJ, Vargas F, Vazquez-Baeza Y, Vogtmann E, von Hippel M, Walters W, Wan Y, Wang M, Warren J, Weber KC, Williamson CHD, Willis AD, Xu ZZ, Zaneveld JR, Zhang Y, Zhu Q, Knight R & Caporaso JG (2019) Reproducible, interactive, scalable and extensible microbiome data science using QIIME 2. Nat Biotechnol 37, 852–857, doi: 10.1038/s41587-019-0209-9.

